# Redox activation of ATG5 licenses autophagy upon nutrient restriction

**DOI:** 10.64898/2026.09.23.753803

**Authors:** Sanghee Shin, Ekaterina D. Korobkina, Claudia Lennicke, Nils Burger, Bingsen Zhang, Haopeng Xiao, Yaoyu Wang, Jonathan J. Petrocelli, Shelley M. Wei, David Wood, Yu Lei, Xiaomei Zhang, Maria C. Perez-Matos, Muneeb A. Sharif, Jiuchun Zhang, Helena M. Cochemé, William B. Mair, Edward T. Chouchani

**Affiliations:** Department of Cancer Biology, Dana–Farber Cancer Institute, Boston, MA, USA; Department of Cell Biology, Harvard Medical School, Boston, MA, USA; MRC Laboratory of Medical Science (LMS), London, UK; Institute of Clinical Sciences, Imperial College London, Hammersmith Hospital Campus, London, UK; Department of Biochemistry, Stanford University School of Medicine, Stanford, CA, USA; Stanford Cancer Institute, Stanford University School of Medicine, Stanford, CA, USA; Department of Molecular Metabolism, Harvard T.H. Chan School of Public Health, Boston, MA, USA; Howard Hughes Medical Institute, Chevy Chase, MD, USA

## Abstract

Dietary restriction (DR) protects against metabolic disease, extends lifespan, and is associated with remodeling of tissue reactive oxygen species (ROS). ROS control biological adaptation through reversible oxidation of protein cysteines, yet the targets of DR-initiated redox signaling are unknown. Here we generate OxiDR, a tissue-resolved atlas of the cysteine redox proteome that quantifies oxidation state under DR. Rather than oxidizing the proteome broadly, DR selectively targets a high-amplitude set of cysteines in a tissue-specific manner, allowing systematic classification of biological processes subject to DR-mediated redox regulation. Among the cysteines most highly oxidized upon DR is Cys19 of the core autophagy protein ATG5. We show oxidation of Cys19 is required for ATG5-mediated autophagosome formation and for autophagy triggered by nutrient restriction in human cells and mice. Reversible oxidation of this cysteine promotes ATG5 binding to ATG10, thus forming the ATG5–ATG12 conjugate that lipidates LC3B/ATG8 and matures the autophagosome. In mice, loss of this redox switch prevents effective initiation of autophagy upon nutrient restriction, resulting in gross tissue pathology and rapid onset of mortality. The autophagic response to nutrient restriction is thus gated by oxidation of a single cysteine.

---

Dietary restriction (DR), the reduction of nutrient intake without malnutrition, has been extensively studied for its effects on metabolic health and healthy lifespan across a wide range of organisms, from yeast to mammals (*1–5*). Underlying DR are mechanisms that have been conserved throughout evolution to optimize survival in times of limited food availability. Studies across model organisms have consistently shown that DR extends lifespan and protects against a myriad of metabolic diseases (*6–9*). Understanding the evolutionarily conserved mechanisms of DR’s impact on metabolic health and lifespan holds promise for unraveling the fundamental processes that govern diseases of aging and developing interventions to promote metabolic health in humans.

A conserved metabolic response to DR is remodeling of cellular redox metabolism (*10–12*). These alterations regulate a mode of post-translational signaling: the generation of reactive oxygen species (ROS) and related species that alter protein function through covalent modification of cysteine residues (*13, 14*). Due to the rapid and reversible nature of cysteine oxidation, it is used to modulate protein function and localization (*15–17*). Consequently, redox modification of protein cysteines is implicated in a wide range of regulatory processes. Furthermore, alterations in ROS and redox signaling are among the central phenomena associated with age-related pathophysiology as well as interventions that improve longevity (*12, 18–20*).

The protein modifications that provide the mechanistic basis for redox regulation upon DR remain unknown. Here, we apply a cysteine derivatization and enrichment method coupled with multiplexed proteomics to comprehensively map tissue-specific protein cysteine oxidation *in vivo* upon DR in aging mice. This OxiDR dataset quantifies the % reversible modification of over 171,000 cysteine sites across six mouse tissues in young and old mice under *ad libitum* (AL) feeding conditions and upon DR, corresponding to ∼37,000 unique sites across ∼10,000 proteins. This landscape represents the first quantitative analysis of the mouse cysteine redox proteome regulated by DR.

From this dataset, we observe a highly selective remodeling of cysteine oxidation state on proteins across six tissues, with many of these targets known to be relevant to metabolic disease and diseases of aging. We systematically classify these proteins to define biological processes that underlie aging and longevity that are subject to redox regulation. Among the cysteines that undergo the most pronounced redox modification induced by DR is Cys19 on the pivotal autophagy protein ATG5. We show that reversible oxidation of this cysteine plays a critical role in ATG5-mediated autophagosome formation and autophagy upon nutrient restriction. Mechanistically, oxidation of Cys19 is required for ATG5 interaction with ATG10, which facilitates the transfer of ATG5 to ATG12 and formation of the ATG5-ATG12 complex. Cells lacking Cys19 on ATG5 exhibit profound defects in autophagy due to the inability of ATG5 to form a complex with ATG12 and initiate LC3B lipidation. In mice, loss of this redox switch prevents initiation of autophagy upon nutrient restriction and drives rapid morbidity and mortality upon fasting. Together, we describe a redox activation mechanism for effective induction of autophagy upon nutrient restriction. More generally, these findings provide a comprehensive analysis of redox-signaling networks in living tissues, which can be accessed through an interactive web resource at http://oxidr-lb-260235877.us-east-1.elb.amazonaws.com/OxiDR/

## RESULTS

### A Quantitative Tissue-Specific Landscape of Protein Cysteine Oxidation in Mice Upon DR

Despite the apparent importance of redox signaling in DR, there remains a lack of comprehensive information on the specific protein modifications underlying this form of biological regulation. We recently developed a method for cysteine derivatization and enrichment coupled with multiplexed proteomic mass spectrometry (MS), termed CPT-MS. CPT-MS provides the basis for comprehensive characterization of protein cysteine oxidation *in vivo*, allowing for simultaneous quantification of the reversible oxidation state for tens of thousands of protein cysteine residues in a single experiment (*21, 22*). Here, we used this approach to systematically define the landscape of protein cysteine targets of redox regulation upon DR in living mice **(Fig. 1A)**. We deployed a labeling method to enable comprehensive quantification of reversibly oxidized cysteine thiols throughout the proteome. This approach was complemented by Tandem Mass Tag (TMT) multiplexing, facilitating simultaneous analysis of eight biological replicates within a single experiment **(Fig. 1B)**. Using this strategy, the percentage of reversible cysteine redox modification can be determined, accounting for variations in protein abundance (*21*).

**Figure 1.**
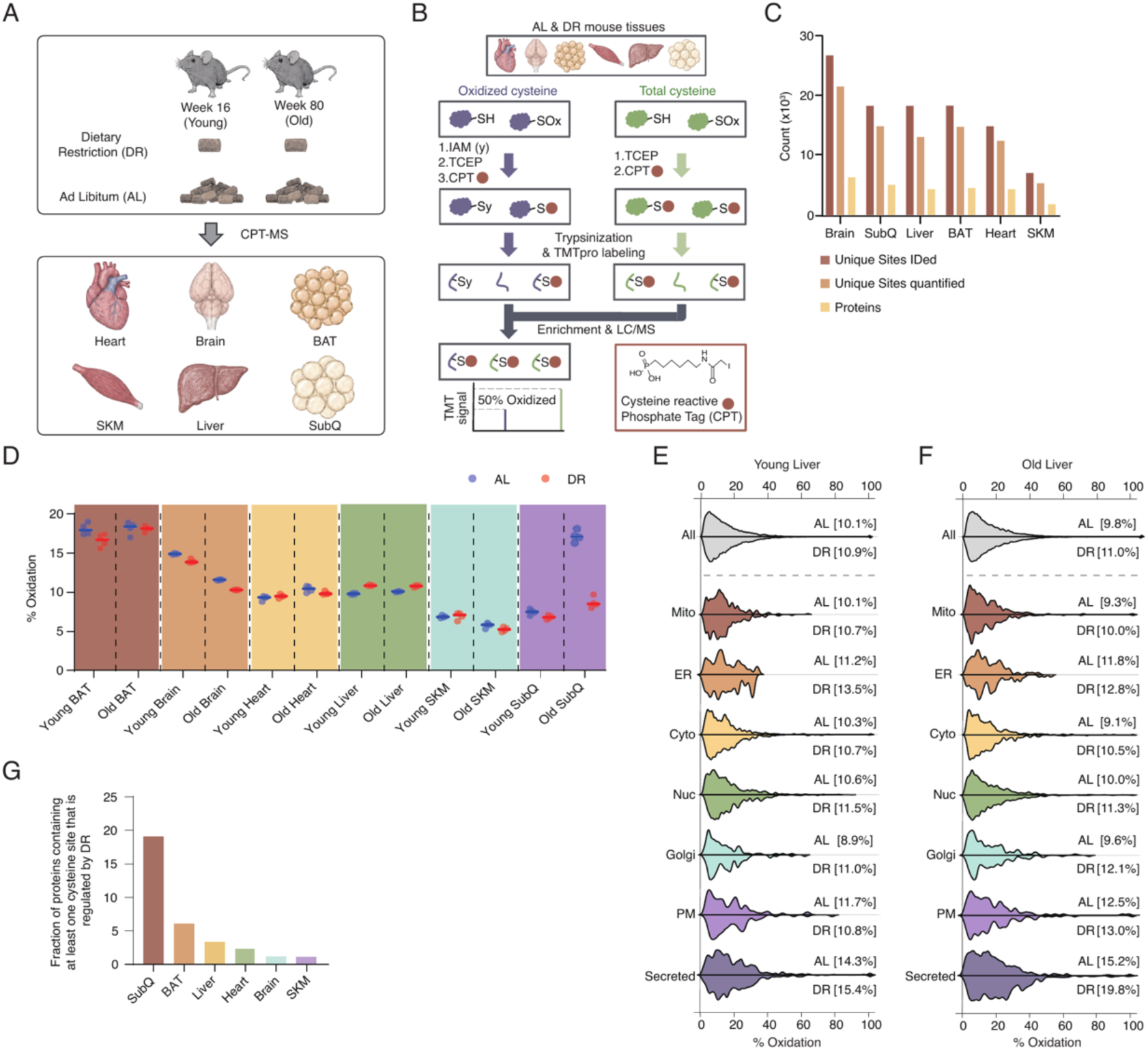
The quantitative tissue-specific cysteine redox proteome regulated by DR. **A.** Study overview of 6 tissues harvested from young and old mice that are subjected to either Dietary restriction (DR) or Ad libitum (AL) feeding. BAT, brown adipose tissue; SKM, skeletal muscle; SubQ, subcutaneous fat. **B.** Schematic overview of CPT-MS proteomics to determine % reversible modification of the cysteine proteome. **C.** Number of cysteines identified and quantified in each tissue. n = 4 mice per group. **D.** Distribution of % cysteine modification for each tissue under different conditions. Solid lines indicate median for each group. n = 4 mice per group. **E.** Percent oxidation distribution of cysteine proteome across subcellular compartments in liver of young AL fed and DR mice. Median percent oxidation values for each group are shown. n = 4 mice per group. **F.** Percent oxidation distribution of cysteine proteome across subcellular compartments in liver of old AL fed and DR mice. n = 4 mice per group. **G.** Fraction of proteins containing at least one cysteine site that is regulated by DR across different tissues. n = 4 mice per group.

To determine cysteine oxidation sites regulated by DR in living tissues, six organs were harvested from male C57BL/6J mice at 16 weeks and 80 weeks of age on a lifelong AL diet or subjected to DR. We used a DR protocol that is well established to initiate lifespan extension and healthspan benefits (*23–26*). DR was initiated at 14 weeks of age at 10% restriction of calories compared to AL, increasing to 25% at 15 weeks, then further to 40% at 16 weeks, which was maintained for the remainder of the experiment. Four biological replicates were harvested for each tissue at each age and diet, then subjected to the CPT redox proteomics workflow **(Fig. 1B)**. The final dataset, termed OxiDR, contained over 171,000 individual cysteine sites mapped across all tissues **(Fig. 1C and table S1)**, corresponding to approximately 37,000 unique cysteine sites on 10,000 proteins.

### Population Characteristics of Redox-Regulated Proteins Upon DR

Consistent with our previous study, we found that the overall cellular and compartmentalized redox tone of most tissues was similar when comparing young and old AL-fed mice, supporting the notion that aging does not generally coincide with bulk shifts in oxidation of protein cysteine residues (*21*). Interestingly, in most tissues, DR did not drive substantial changes in bulk protein cysteine oxidation **(Fig. 1D)**. Moreover, at the organellar level, DR did not significantly alter the bulk subcellular redox tone **(Fig. 1E,F and fig. S1A)**. One exception to this was found in subcutaneous adipose tissue (SubQ), for which we observed a significant decrease in bulk protein cysteine oxidation upon DR in 80-week-old mice, which was attributable to a decrease in bulk redox tone in numerous cellular compartments including mitochondria, ER and secreted proteins **(Fig. 1D and fig. S1A)**. We next performed protein- and site-level analysis to define targets that are redox regulated upon DR across mouse tissues. In all tissues, most protein cysteine residues exhibited no DR-dependent change in redox modification **(Fig. 1G)**. Nevertheless, each tissue had a sub-population of cysteines that were regulated by DR (>15% change in modification state). Across the dataset, 2,052 cysteine residues were differentially regulated in the context of DR in at least one tissue (5.6% of the entire mapped population). Because SubQ exhibited a bulk oxidation shift across multiple cellular compartments upon DR, its inclusion likely inflates the regulated population with cysteines responding to a global change in tissue redox poise rather than targeted modification. Excluding SubQ tissue from this analysis, where bulk oxidation shifts were observed, we found that 839 cysteine residues were differentially regulated in the context of DR in at least one tissue (2.3% of the entire mapped population). By excluding SubQ, this more conservative set of cysteines is more likely to reflect site-selective redox signaling than broader changes in the cellular redox environment.

Further, we found there was no correlation between protein abundance and extent of cysteine modification upon DR in any tissue **(fig. S1B)**. Instead, examination of predicted structural elements within proteins revealed that highly modified cysteine sites were associated with differences in local sequence features. Consistent with prior findings (*21*), motif analysis revealed enriched patterns of positively charged residues flanking redox-sensitive cysteines. As shown previously (*21*), these local electrostatic environments could modulate the thiol-thiolate equilibrium, thereby tuning redox reactivity at specific sites. Overall, these data suggest that cysteine responsiveness to redox changes upon DR is encoded in part by the proximal amino acid environment **(fig. S1C)**.

### DR-mediated Redox Regulation of Individual Proteins Is Tissue Specific

DR induced negligible population-level changes to the redox proteome in most tissues (**Fig. 1D**). We instead observed a substantial number of individual cysteines that exhibited selective dynamic modification in each tissue upon DR **(Fig. 2A and table S2)**. Overall, DR-regulated cysteines exhibited marked tissue specificity, with DR-driven modifications often distinct to each tissue **(Fig. 2B-D)**. The correlation in modification states of individual DR-regulated cysteine residues between different tissues was notably weak **(Fig. 2E,F)**. Furthermore, when performing pairwise comparisons of individual tissues with similar overall redox profiles, consistent clusters of highly regulated cysteine sites initiated by DR emerged, which are specific to each tissue **(Fig. 2B,C)**. Remarkably, many of these tissue-specific DR-regulated sites are located on proteins that are expressed in all tissues **(fig. S2A,B)**. We analyzed each cysteine residue within the OxiDR dataset to evaluate its dynamic oxidation driven by DR (**table. S2**), revealing that most DR-regulated sites displayed a high degree of tissue-selective modification **(Fig. 2D-H)**. These findings offer evidence that broadly expressed proteins are subject to dynamic redox regulation upon DR that is distinct in each tissue, likely arising from localized upstream redox metabolic processes that respond to DR differently depending on the tissue. In support of this notion, we observed that the expression levels of major enzymes involved in the metabolism of ROS and enzymes responsible for regulating subcellular thiol redox state also exhibit DR-regulated abundance **(fig. S2C-F)**. Similarly, established protein sources of ROS also display abundance profiles that are regulated by DR **(fig. S2G-J)**.

**Figure 2.**
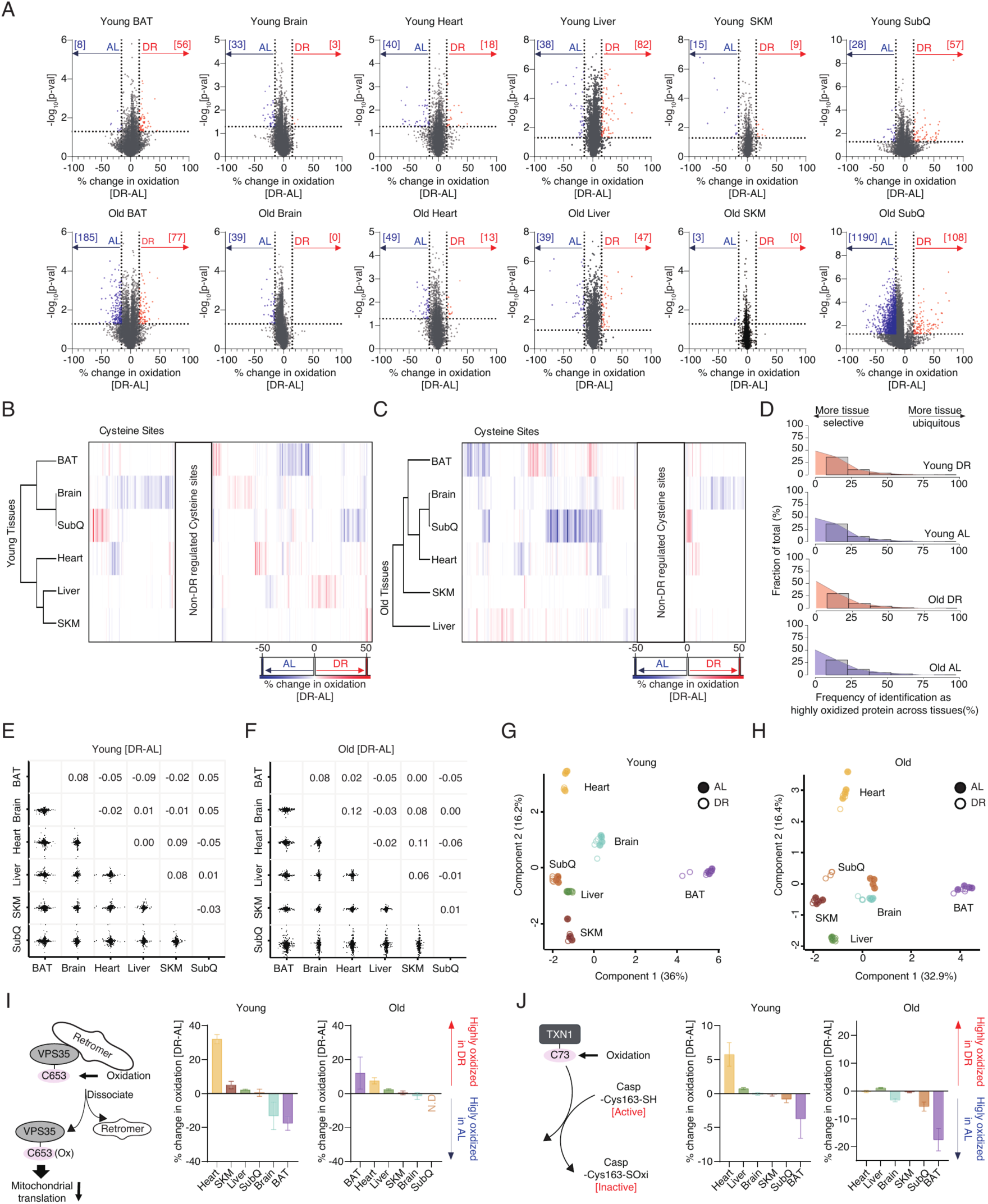
Mapping tissue-specific redox regulation of individual cysteine sites upon DR. **A.** Differential cysteine modification state of individual cysteines across the mapped cysteine proteome comparing DR to AL in each mouse tissue at each age. n = 4 mice per group. **B.** Heatmap cluster of cysteine sites showing concordant change in oxidation state upon DR across all tissues in young mice. n = 4 mice per group. **C.** Heatmap cluster of cysteine sites showing concordant change in oxidation state upon DR across all tissues in old mice. n = 4 mice per group. **D.** Frequency distribution plot of proteins that contain at least one cysteine site >20% differentially oxidized upon DR across tissues. n = 4 mice per group. **E.** Correlation matrix of the percentage change in cysteine oxidation across all sites upon DR across young tissues. n = 4 mice per group. **F.** Correlation matrix of the percentage change in cysteine oxidation across all sites upon DR across old tissues. n = 4 mice per group. **G.** Principal component analysis of % oxidation state of cysteine proteomes from young mouse tissues under AL and DR conditions. n = 4 mice per group. **H.** Principal component analysis of % oxidation state of cysteine proteomes from old mouse tissues under AL and DR conditions. n = 4 mice per group. **I.** Schematic representation and quantification of tissue-specific redox regulation of VPS35 Cys653 upon DR. n = 4 mice per group. **J.** Schematic representation and quantification of tissue-specific redox regulation of TXN1 Cys73 upon DR. n = 4 mice per group. All data are presented as mean ± SD.

We identified numerous examples of tissue-specific regulation of established functional cysteines on ubiquitously expressed proteins initiated by DR. For instance, reversible oxidative modification of Cys653 on VPS35 is known to regulate retromer membrane association, plasma membrane composition, and mitochondrial translation (*27*). Here, we found that tissues exhibit differential redox modification of this regulatory site upon DR, suggesting DR can tune mitochondrial translation and ROS production through regulation of this site **(Fig. 2I)**. Another established example was found in the case of oxidative modification of Cys73 on thioredoxin 1 (TXN1), which can be reversibly transferred to Cys163 on caspase-3 (Casp-3), thereby inhibiting its enzymatic activity (*28*). Here, upon DR, we observed the redox status of this site varied markedly upon DR in a tissue-specific manner. This suggests that different tissues may tune the activity of caspase-3 through modulating the oxidation state of TXN1 Cys73 upon DR **(Fig. 2J)**. These examples highlight how the OxiDR dataset can nominate tissue specific redox regulatory nodes relevant to DR, encompassing both established regulatory sites, and hundreds of previously uncharacterized cysteine sites across the proteome, which are elaborated on below.

### Defining DR-Mediated Redox Regulation of Protein Networks

Many tissue-specific cysteine oxidation sites regulated by DR converge toward distinct biological processes **(fig. S2K)**. We postulated that overlaying DR-regulated cysteine oxidation data onto proteomic networks could reveal coordinated cysteine modifications governing shared biological activities of proteins. We integrated OxiDR with BioPlex 3.0, a comprehensive proteomic interactome (*29*), to elucidate redox-modified protein networks contributing to both tissue-specific and tissue-ubiquitous biological processes regulated by DR. Protein networks displaying coordinated redox regulation of cysteines upon DR, defined by a cysteine oxidation shift exceeding 15%, were categorized as “DR-regulated redox networks” within each tissue **(Fig. 3A)**. This approach enabled us to identify DR-regulated protein redox networks that are present across all tissues and those exhibiting tissue-specific patterns **(fig. S3A-C)**. Excluding SubQ tissue from this analysis, where bulk oxidation shifts were observed, we found among the 1,423 protein networks established in BioPlex 3.0, 217 exhibited coordinated redox regulation of cysteines upon DR in at least one tissue **(Fig. 3A, fig. S3A and table S3)**.

**Figure 3.**
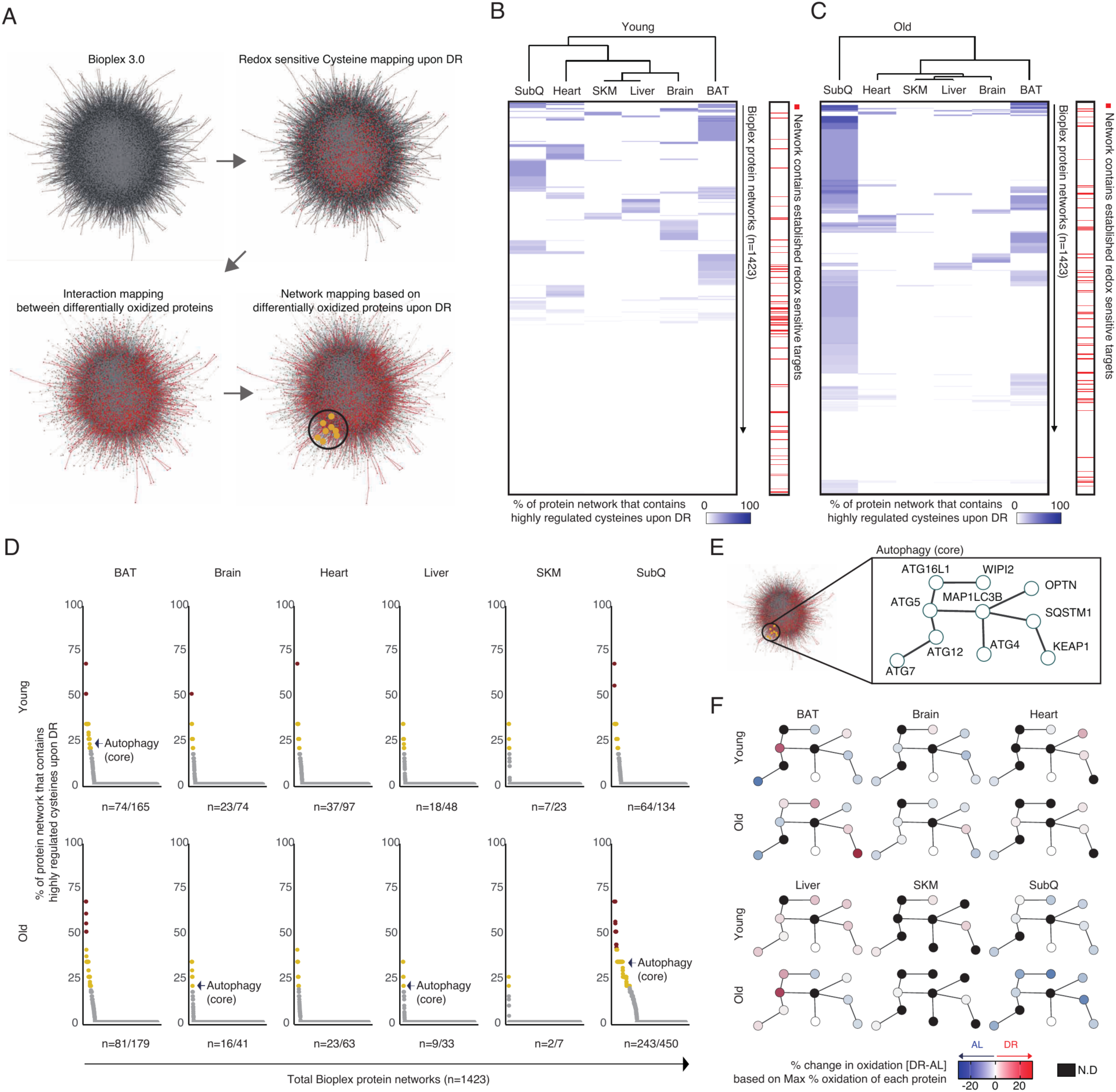
Tissue-specific redox networks regulated by DR. **A.** Visualization of Bioplex 3.0, which includes 129687 interactions (lines) across 13957 proteins (nodes). Cysteine oxidation networks are defined by mapping proteins harboring differentially oxidized cysteines (red node). n = 4 mice per group. **B.** Clustered heatmap showing the proportion of protein redox networks containing cysteines that are regulated upon DR in young mouse tissues, with indications of redox networks that contain previously reported redox sensitive protein targets. n = 4 mice per group. **C.** Clustered heatmap showing the proportion of protein redox networks containing cysteines that are regulated upon DR in old mouse tissues, with indications of redox networks that contain reported redox sensitive protein targets. n = 4 mice per group. **D.** Extent of coordinated redox modification of protein networks across mouse tissues. n = 4 mice per group. **E.** Schematic representation of the protein network involved in autophagy pathway. **F.** Percentage changes in protein oxidation upon DR, corresponding to the network shown in **(E)**. n = 4 mice per group.

Among the proteins present in DR-regulated redox networks (**Fig. 3B,C**), 64 included protein cysteines that had been previously reported as susceptible to oxidative modifications in various biological models, including yeast, flies and mice (*30–32*). Characteristically, nearly all these redox networks displayed some degree of tissue specificity (**Fig. 3B,C and fig. S3B)**. Redox networks nominated numerous coordinated processes subject to redox regulation upon DR. Among the most prominent DR-regulated redox networks contained proteins that control autophagosome formation and autophagy, a fundamental cellular process tightly linked to longevity and healthspan (*33, 34*) **(Fig. 3D-F)**. We explored this finding in more detail in a subsequent section.

### DR-Dependent Redox Regulation of Longevity-associated proteins

Our above analysis enumerated proteins and protein networks subject to coordinated cysteine oxidation upon DR. Many of these proteins are involved in cellular processes and have been associated with interventions that extend lifespan, implying that proteins influenced by DR-dependent redox regulation could serve as functional targets contributing to the longevity effects of DR. To explore this phenomenon in more detail we mapped DR-dependent redox targets onto GenAge, a database containing proteins for which genetic evidence exists for a role in lifespan regulation (*35*). In doing so, we identified several DR-driven redox-regulated sites on well-established longevity targets **(fig. S4A)**. We categorized redox-regulated proteins that exhibited an increase in modification upon DR as “DR-driven redox targets”. Conversely, we classified targets showing a loss of redox modification coinciding with DR as “AL-driven redox targets” **(Fig. 4A)**. Prominent candidates of DR-driven longevity targets were found in the heart, liver and SubQ adipose tissue **(Fig. 4A and fig. S4B,C)**. Intriguingly, AL driven targets were predominantly mitochondrial proteins associated with lipid metabolism, whereas DR-driven sites were associated with a broader range of cellular processes **(fig. S4D)**. Collectively, these findings provide a compendium for understanding mechanisms of DR-driven redox regulation over proteins relevant to longevity. Since many benefits of DR are conserved across evolution, we additionally noted which of these sites exhibited conservation across species. Of the established longevity related proteins, three contained redox regulated cysteines that are conserved across species from *Homo sapiens* to *Drosophila melanogaster*: ATG5 Cys19, TPP2 Cys733, and COQ7 Cys137 **(Fig. 4A)**.

**Figure 4.**
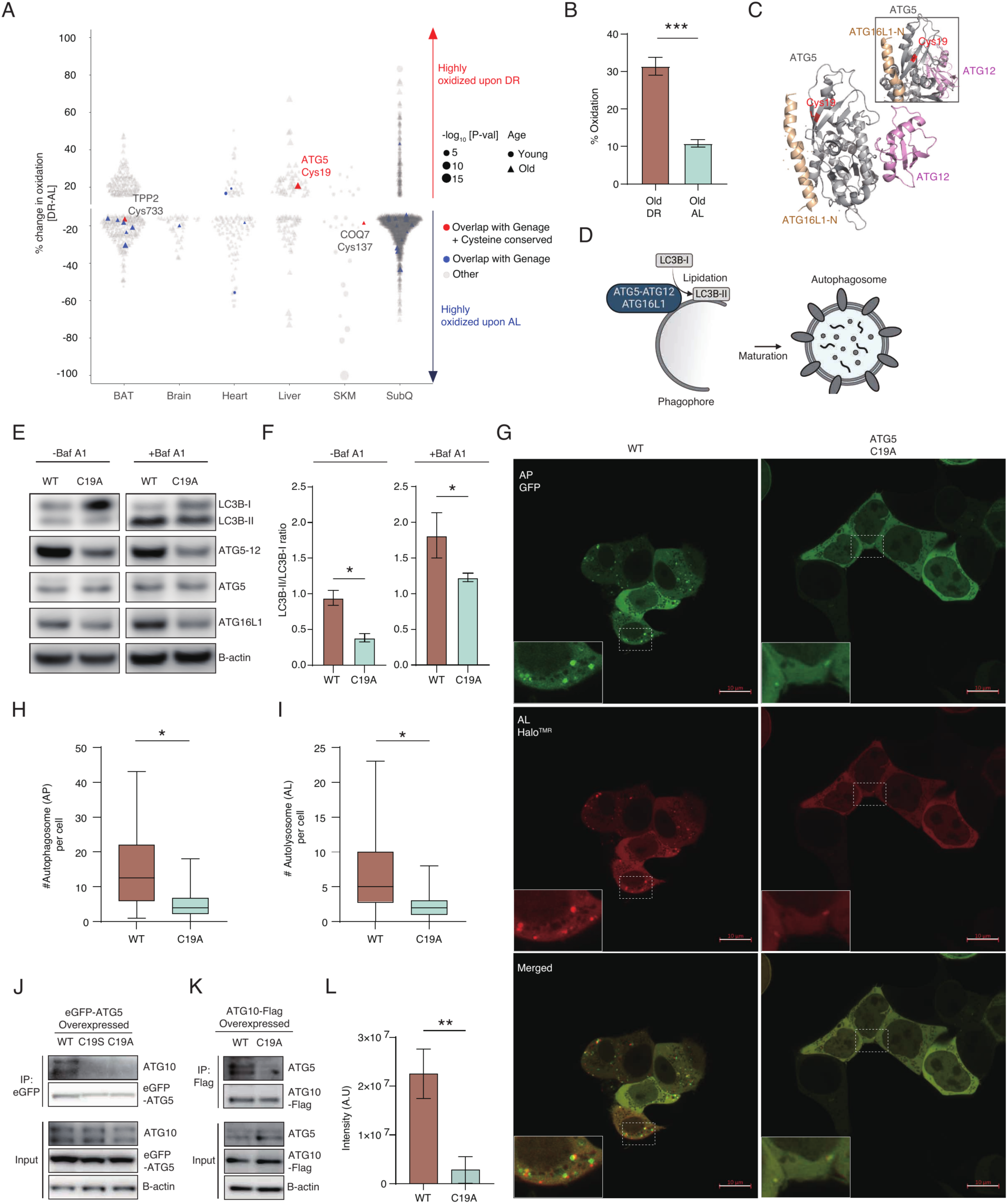
Redox regulation of ATG5 Cys19 modulates autophagosome formation and is required for fasting induced autophagy. **A.** Scatter plot of cysteine redox sites across tissues that are significantly regulated by DR. Red dots indicate proteins overlapping with the GenAge database which contain conserved regulated cysteine sites across species (from mammals to *D. melanogaster*). Blue dots indicate proteins overlapping with GenAge database. **B.** Oxidation levels of ATG5 Cys19 mapped in old mouse liver. n = 4 mice per group. **C.** Crystal structure of the ATG5-ATG12 and ATG16L1 N-terminal fragment complex [PDB: 4GDK] with an inset showing an enlarged view of the ATG5 Cys19 region (highlighted in red). **D.** Schematic of LC3B lipidation mediated by the ATG5-12-16L1 complex during autophagosome elongation and maturation. **E.** Immunoblot of LC3B, ATG5-12, and ATG16L1 from WT and ATG5 C19A cells, with or without Bafilomycin A1 pretreatment (10 nM, 2hr). C19A cells exhibit significantly reduced LC3B lipidation along with decreased ATG5-12 covalent complex and lower ATG16L1 levels. **F.** LC3B-II (lipidated) to LC3B-I (non-lipidated) ratio measured by immunoblot under basal conditions with or without Bafilomycin A1 pretreatment (10 nM, 2hr). n = 3 cell replicates per group. **G.** Immunofluorescent imaging of autophagic flux in WT and C19A cells using Halo-GFP-LC3B construct. See methods for details of analysis. **H.** Quantification of autophagosomes per cell corresponding to experiment in **(G)**. ATG5 C19A cells show significantly lower autophagosome formation. n= 65-78 cells per group. **I.** Quantification of autophagosome per cell corresponding to experiment in **(G)**. ATG5 C19A cells show significantly lower autolysosome formation. n = 65-78 cells per group. **J.** Immunoprecipitation of WT, C19S and C19A eGFP-ATG5 showing interaction of ATG10 with WT ATG5 that is lost in mutant cells. **K.** Immunoprecipitation of ATG10-Flag in either WT or ATG5 C19A cells. WT ATG5 interaction with ATG10 is lost in ATG5 C19A cells. **L.** Densitometry quantification of ATG5 band shown in **(K)**. (A.U = arbitrary unit) n = 3 cell replicates. p-value<0.05 is marked as “*”, p value<0.01 as “**” and p value<0.001 as “***”. Two-tailed Student’s t-test were used for pairwise comparisons. All bar plot data are presented as mean ± SD.

### OxiDR Uncovers a Redox Switch that Regulates Autophagosome Formation

OxiDR analysis of longevity-associated proteins revealed Cys19 on the core autophagy protein ATG5 among the most highly redox-modified cysteines induced by DR in the liver **(Fig. 4A,B)**. ATG5 is widely expressed and particularly abundant in the adrenal gland and liver (*36*) (**fig. S4E**), and plays an essential role in the formation and expansion of the autophagosome(*37, 38*). Autophagy is known to decline with age, whereas ectopic overexpression of ATG5 in mice is sufficient to significantly increase lifespan (*39*). Conversely, KO of ATG5 leads to neonatal lethality in mice (*39–41*). More generally, initiation of autophagy is known be regulated by ROS (*42*), and is critical for the protective and lifespan extending effects of DR (*33, 34, 43, 44*).

While ATG5 exhibited redox modification upon DR **(Fig. 4B)**, we observed no detectable changes in ATG5 protein abundance across diet conditions or during aging **(fig. S4F)**. ATG5 is composed of three domains, including two ubiquitin-like (Ubl) domains (UblA & UblB) connected by an alpha helical bundle where ATG12 is known to conjugate **(Fig. 4C and fig. S4G)**. ATG5 Cys19 is situated within UblA near the established interfaces with other ATG proteins that interact with ATG5 to regulate autophagosome formation (*45–47*) **(Fig. 4C,D)**. On this basis, we examined whether oxidation of Cys19 regulates ATG5 function, and ATG5-dependent mechanisms of autophagy.

We first explored whether oxidation of ATG5 Cys19 induced by DR *in vivo* could be recapitulated in cellular models of nutrient restriction wherein, like DR, autophagy is robustly activated. We examined mouse AML12 cells under acute nutrient restriction that is known to initiate autophagy (*48*). Restriction of nutrients drove a robust and selective increase in ATG5 Cys19 oxidation **(fig. S4H)** to a similar extent as observed in mouse liver upon DR **(Fig. 4B)**. We next tested whether oxidation of Cys19 is associated with autophagosome formation and autophagy.

The proximal step in autophagosome formation initiated by ATG5 requires its localization to lipid membranes where, in complex with ATG16L1 and ATG12, it facilitates lipidation of LC3B (*45, 47, 49*) **(Fig. 4D)**. Notably, preventing protein thiol oxidation with the thiol reducing agent N-acetyl cysteine (NAC) resulted in substantial decrease in a lipidated form of LC3B (LC3B-II) upon nutrient restriction **(fig. S4I)**. Moreover, NAC treatment resulted in a concentration dependent decrease in the abundance of the ATG5-12 conjugate complex **(fig. S4J)**.

We next determined whether the effects of manipulating thiol redox state on ATG5-dependent processes were through regulation of Cys19. We generated cells expressing RFP-tagged LC3B and either wild type (WT) or mutant ATG5 (C19S & C19A) with GFP tagged to the N-terminus. Remarkably, ATG5 C19S & C19A cells exhibited an almost complete abrogation of LC3B puncta in both baseline and nutrient restricted hepatocytes **(fig. S5A-C)**. To examine the above effects using endogenous ATG5 protein, we engineered HEK293T cells where Cys19 on ATG5 was replaced with an alanine **(fig. S6A)**. Again, endogenous ATG5 C19A cells exhibited a significant decrease in lipidation of LC3B **(Fig. 4E,F and fig. S6B-E)**. Next, we monitored autophagic flux using the pulse-chase HaloTag reporter (*50*) **(fig. S6f)**. Loss of ATG5 Cys19 led to a significantly lower rate of autophagosome formation and a resultant approximately two-fold decrease in autophagosome and autolysosome formation **(Fig. 4G-I)**. Together, these findings indicate that the redox regulated ATG5 Cys19 is critical for ATG5-mediated autophagosome formation and autophagy.

### ATG5 Cys19 Mediates Interaction with ATG10 to Form the ATG5-12 complex

We next determined how redox regulated Cys19 controls autophagosome formation. Conjugation of ATG5 to ATG12 is an early step in the assembly of the ATG5-12-16 complex that facilitates autophagosome formation (*51–54*). We found that mutation of ATG5 Cys19 resulted in lower abundance of the ATG5-12 complex under basal conditions and upon nutrient restriction **(Fig. 4E and fig. S6B)**. We first sought to understand how redox modification of ATG5 Cys19 regulates formation of the ATG5-12 complex. Formation of the ATG5-12 complex is facilitated by the E2-like enzyme ATG10, which interacts directly with ATG5 (*55, 56*) **(fig. S6G)**. We examined whether ATG5 interaction with ATG10 required Cys19 by expressing WT, C19S, and C19A ATG5 and performing co-immunoprecipitation (Co-IP) with either endogenous or overexpressed ATG10. Remarkably, we found that ATG10 interaction with ATG5 was lost upon mutation of ATG5 Cys19 **(Fig. 4J-L)**.

To better understand how ATG5 Cys19 mediated interaction with ATG10, we examined the sequence and structural properties of both proteins. We noted that human ATG10 possesses an unstructured region composed of approximately 40 amino acids that is conserved across mammals and contains a conserved cysteine residue at position 68 **(Fig. 5A, fig. S7A and table S4)**, while Cys19 on ATG5 is also highly conserved across mammals **(fig. S7B)**. Based on this pattern of shared conservation, we hypothesized that these two cysteines may form a disulfide to mediate interaction between ATG5 and ATG10 **(Fig. 5B)**. This notion was supported by AlphaFold3 co-fold modeling of the ATG10 Cys68 containing region with ATG5, which predicted direct interaction of these domains with distance between ATG5 Cys19 and ATG10 Cys68 of 2.6 Å **(Fig. 5C)**. To test this hypothesis, we generated several ATG10 mutants, either removing the unstructured loop (LD1 and LD2) or mutating Cys68 to alanine (C68A) **(Fig. 5D)**. Next, we co-transfected ATG10 constructs with either WT or C19A ATG5, followed by Co-IP. These analyses demonstrated that both ATG5 Cys19 and ATG10 Cys68 were required for the two proteins to interact. Loss of Cys68, or the entire unstructured region of ATG10 resulted in complete loss of interaction between the two proteins **(Fig. 5E)**. Likewise, ATG10-pulldown in cells expressing either WT or C19A ATG5 showed that only endogenous WT ATG5 interacted with WT ATG10 **(fig. S7C)**. Based on these findings we conclude that oxidative modification of ATG5 Cys19 to form an interprotein disulfide with ATG10 Cys68 promotes the ATG5-ATG10 interaction. Because ATG10 is needed to load ATG12 on ATG5, this disulfide interaction is likely key for facilitating ATG5-ATG12 complex formation.

**Figure 5.**
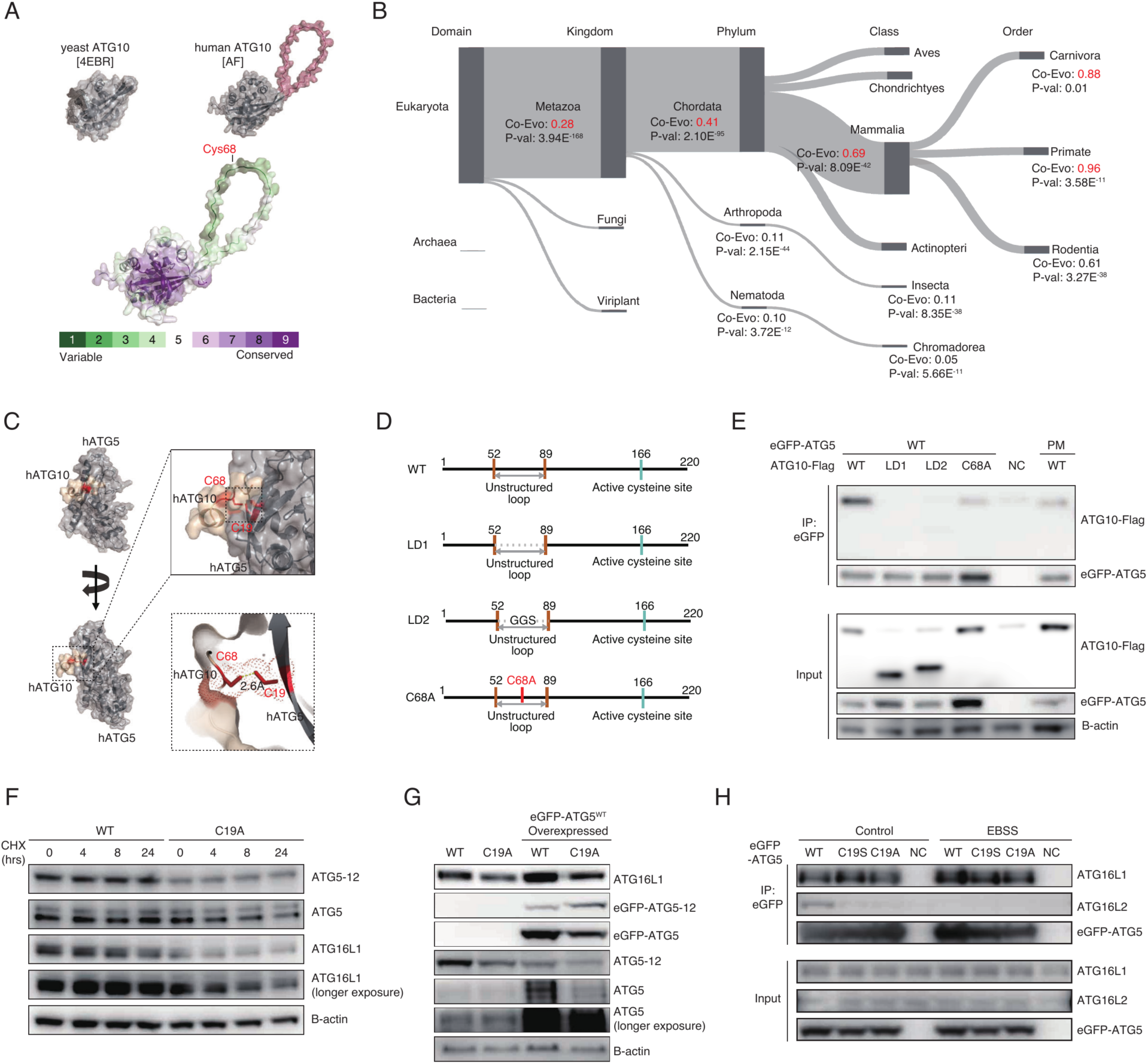
ATG5 Cys19 mediates interaction with ATG10 to facilitate ATG5-12 complex formation. **A.** Crystal structure of yeast ATG10 (top left, 4EBR), AlphaFold3 predicted structure of human ATG10 (top right), and evolutionary amino acid conservation analysis of human ATG10 performed via Consurf (bottom). **B.** Conservation analysis of ATG5 Cys19 and ATG10 Cys68 among diverse species in the OMA database, suggesting a shared pattern of conservation between these two independent cysteine residues (Co-Evo score ranges from 0 to 1). **C.** AlphaFold3 co-fold analysis of the complex between humans ATG5 (hATG5) and the human ATG10 (hATG10) fragment containing Cys68. **D.** Schematic of ATG10 constructs used to test ATG10 Cys68 dependent interaction with ATG5. **E.** ATG5 immunoprecipitation with overexpressed ATG10 constructs shown in **(D)** in the presence of either WT or C19A ATG5. ATG10 Cys68 and its surrounding unstructured region are required for interaction with ATG5. **F.** Assessment of ATG5 and ATG16L1 posttranslational stability using cycloheximide (20 µg/ml) in WT and ATG5 C19A cells. **G.** Assessment of ATG16L1 abundance upon overexpression of WT ATG5 in ATG5 C19A cells. **H.** Immunoprecipitation of overexpressed WT, C19S and C19A eGFP-ATG5 showing selective interaction of ATG16L2 with WT ATG5 under basal condition.

### Redox Driven Conjugation of ATG5 to ATG12 protects ATG5 from Proteasomal Degradation

ATG5 is protected from degradation when it forms a complex of ATG5-12-16L1 or ATG5-12-16L2, while free ATG5 and ATG16L1 exhibit a shorter half-life due to enhanced proteasomal degradation **(fig. S6G)** (*57*). We therefore examined post-translational stability of these factors upon loss of redox regulated ATG5 Cys19. We found that loss of Cys19 on ATG5 resulted in reduced half-life of endogenous ATG5 and ATG16L1 but not the covalently conjugated ATG5-12 complex **(Fig. 5F)**. In ATG5 C19A cells, the rapid degradation of unbound ATG5 and ATG16L1 was inhibited by the proteasome inhibitor MG132 **(fig. S7D)**. Intriguingly, the generation of the apoptosis-inducing truncated form of ATG5 (tATG5) (*58*) was also dependent on Cys19 **(fig. S7D)**, suggesting this residue might govern susceptibility to proteolytic cleavage. Collectively, we observed that the mutation of Cys19 on ATG5 impairs the formation of ATG5-12 complex, leading to elevated levels of unbound ATG5 and ATG16L1, which are preferentially targeted for proteasomal degradation. Notably, the reduced level of ATG16L1 was rescued by overexpressing WT ATG5 in ATG5 C19A cells **(Fig. 5G)**. Additionally, we examined ATG16L2, a paralog of the well-characterized ATG16L1 and a known ATG5 interacting protein that stabilizes ATG5. Co-IP revealed that WT, C19S and C19A ATG5 associate with ATG16L1 at comparable levels. However, WT ATG5 exhibited a notably stronger interaction with ATG16L2 relative to the C19S and C19A variants **(Fig. 5H)**. Although the role of ATG16L2 in autophagy remains poorly understood, it is known to bind ATG5 and prevent its proteasomal degradation (*57*).

### ATG5 Cys19 is required for autophagy and survival upon fasting *in vivo*

The above results establish ATG5 Cys19 as a critical redox switch for induction of autophagy upon nutrient restriction. We next tested whether this mechanism is required for the protective physiology observed upon nutrient restriction *in vivo*. To do so, we generated ATG5 C19A point mutant mice. C19A homozygotes were born at Mendelian ratios and appeared grossly normal, with no difference in body weight or growth compared to WT littermates under AL feeding conditions **(fig. S8A,B)**. Baseline analyses revealed a subtle metabolic phenotype whereby both female and male C19A mice exhibited lower resting oxygen consumption and energy expenditure than WT, indicating a modest systemic reduction in metabolic rate **(Fig. 6A and fig. S8C)**. Notably, female C19A mice exhibited significantly higher liver weight than WT, reminiscent of hepatic enlargement seen in other autophagy-deficient genetic models (*59*). No obvious gross changes were seen in other organs at baseline **(fig. S8D)**.

**Figure 6.**
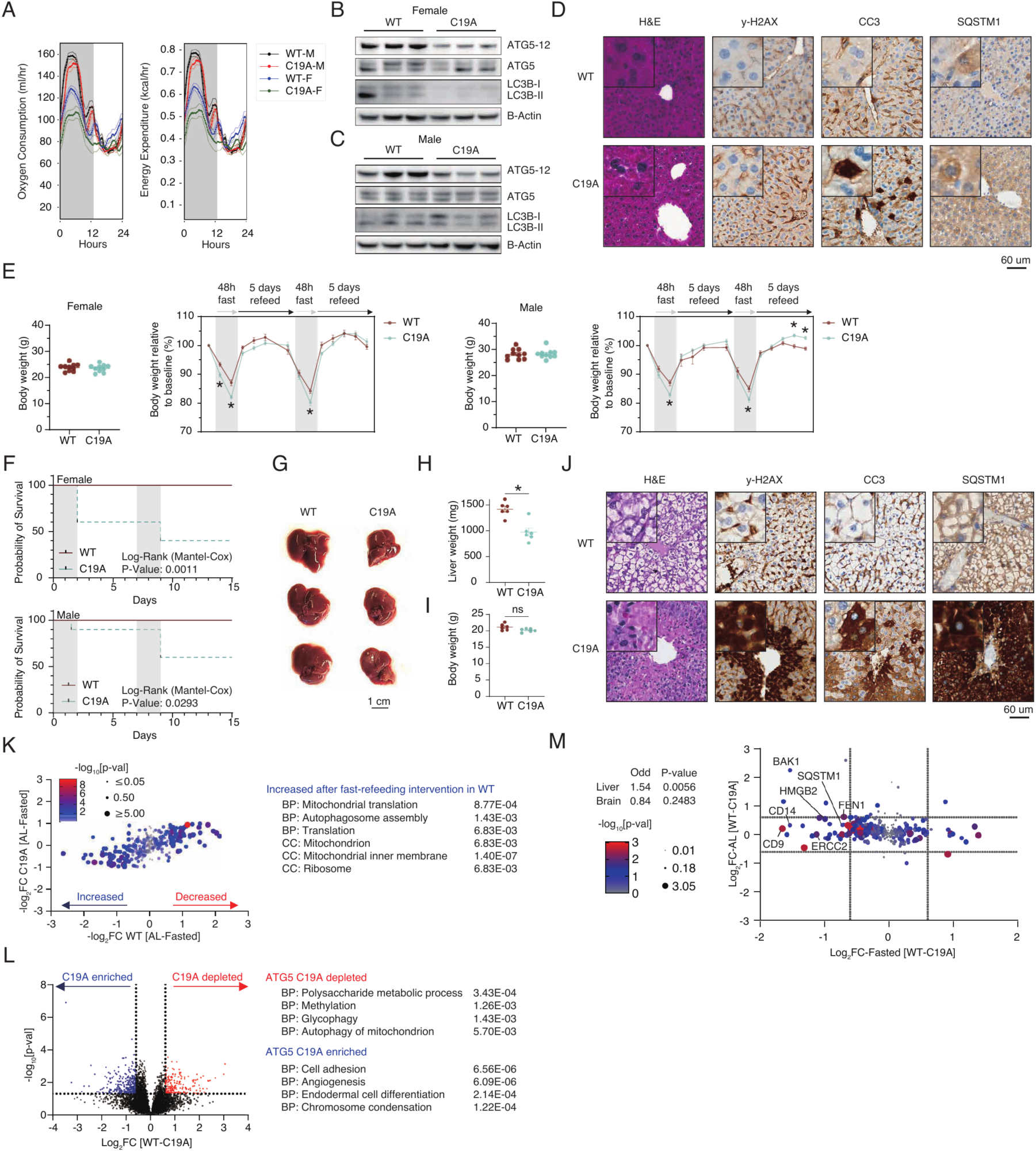
ATG5 Cys19 is required for the physiologic fasting response. **A.** Measurement of oxygen consumption (ml/hr) and energy expenditure (Kcal/hr) in WT and ATG5 C19A mice. n = 6 mice **B.** Immunoblot of WT and ATG5 C19A female mice showing reduced ATG5-12 complex. **C.** Immunoblot of WT and ATG5 C19A male mice showing reduced ATG5-12 complex. **D.** Hematoxylin and Eosin (H&E) staining and immunohistochemical (IHC) staining of y-H2AX, cleaved caspase-3 (CC3) and SQSTM1/p62 from WT and ATG5 C19A mice livers under AL feeding conditions. **E.** Schematic of 2 week intervention of 48 hours intermittent fasting, along with body weight traces of both female and male mice. n = 10 mice per group. **F.** Survival of mice during the intermittent fasting intervention showing that ATG5 C19A mice are intolerant to acute fasting. n = 10 mice per group. **G.** Livers from WT and ATG5 C19A female mice after fasting and 24-hour refeeding. **H.** Liver mass measurements corresponding to **(G)**. n = 6 mice per group. **I.** Bodyweight measurement of female mice, corresponding to **(G)**. n = 6 mice per group. **J.** Hematoxylin and Eosin (H&E) staining and immunohistochemical (IHC) staining of y-H2AX, cleaved caspase-3 (CC3) and SQSTM1/p62 from WT and ATG5 C19A mice livers under following 24 hour fast and refeeding. **K.** Proteomic analysis of livers comparing AL and acute fasted conditions, highlighting proteins that are differentially regulated in either WT or ATG5 C19A mice basally or following fasting. n = 5-6 mice per group. **L.** Proteomic analysis of livers comparing WT and ATG5 C19A after fasting and 24-hour refeeding, highlighting proteins that are differentially regulated in WT liver only. n = 5-6 mice per group. **M.** Proteomic analysis of known aging markers in livers from AL and fasted WT and ATG5 C19A mice. n = 5-6 mice per group. p-value<0.05 is marked as “*”. Two-tailed Student’s t-test were used for pairwise comparisons. All data are presented as mean ± SEM.

Histological and molecular analysis of AL-fed mice indicated modest autophagy impairment and cellular stress upon loss of ATG5 Cys19. Liver immunoblotting confirmed that the C19A mutation reduced formation of ATG5-12 conjugate while ATG5 protein was unchanged **(Fig. 6B,C)**, suggesting less autophagosome formation even under unperturbed conditions. We next examined known molecular hallmarks downstream of impaired autophagy. C19A livers and brains showed modest accumulation of the autophagy receptor SQSTM 1/p62 and higher levels of cleaved caspase-3 (CC3) compared to WT **(Fig. 6D and fig. S8E)**. SQSTM1/p62 accumulation is indicative of reduced autophagic flux and its dysregulation is linked to accelerated aging and age-related pathology (*60–62*). Elevated CC3 reflects increased apoptosis, which is known to rise in response to impaired autophagy, and is known to increase in aging tissues (*63, 64*) **(Fig. 6D and fig. S8E)**. Beyond these markers, proteomic profiling under fed conditions showed few genotype differences. Differential protein expression between WT and C19A livers or brains was minimal **(fig. S8F-H)**. These baseline proteomic data indicate that loss of Cys19 on ATG5 causes modest tissue remodeling absent substantial physiological consequences.

Importantly, our data showed that ATG5 Cys19 oxidation is induced by DR and acute nutrient restriction, and is critical for induction of autophagy in this context. On this basis, we monitored the effects of nutrient restriction on mice lacking ATG5 Cys19. To do so we applied a 48hr fasting protocol known to initiate physiological benefits associated with longevity and long-term DR (*65–68*). Gross physiology of WT mice responded as expected without apparent adverse response **(Fig. 6E,F and fig. S8I-K)**. However, over the first two bouts of fasting both female and male ATG5 C19A mice displayed compromised tolerance to nutrient deprivation, characterized by excessive body weight loss **(Fig. 6E and fig. S8K)**, impaired grip strength **(fig. S8I)**, and reduced locomotor activity **(fig. S8J)** relative to WT controls. Furthermore, C19A mice developed severe physiological deterioration during fasting with some animals reaching predefined humane endpoints **(fig. S8K and video S1,2)**. Notably, even after 24 hours of AL refeeding following a single fast, although total body weights were comparable at the end of the intervention, the livers of fasted C19A mice were significantly smaller than those of WT mice **(Fig. 6G-I)**, suggesting impaired hepatocyte proliferation and regeneration. Consistent with this, WT livers showed ballooning-shaped hepatocytes, a morphology associated with rapid hepatocyte regeneration (*69, 70*). Interestingly, ATG5 C19A livers lacked these features **(Fig. 6J)**. Moreover, C19A livers exhibited a dramatic increase in the aging- and stress-associated markers γH2AX (*71, 72*), CC3 (*63, 64*) and SQSTM1/p62 (*60–62*) **(Fig. 6J)**, a stark contrast compared to WT. This striking phenotype demonstrates that mice lacking ATG5 Cys19 cannot tolerate acute nutrient restriction.

To understand the molecular basis of this gross incapacity to tolerate fasting we examined molecular and proteomic changes in WT and ATG5 C19A mice both at baseline and following fasting. As expected, fasting induced extensive proteome remodeling in WT mouse liver and brain **(fig. S8M,N),** characterized by a coordinated reduction of proteins related to lipid and fatty acid metabolism and remodeling of the mitochondrial proteome **(fig. S8N)**, consistent with prior data (*73, 74*). This transition was accompanied by robust upregulation of protein markers of autophagy **(Fig. 6K and fig. S8N)**. Remarkably, C19A livers lacked this proteome remodeling observed in WT mice **(Fig. 6K and fig.S8O)**. Moreover, like C19A livers, C19A brains exhibited a dramatic increase in the aging- and stress-associated markers γH2AX (*71, 72*), CC3 (*63, 64*) and SQSTM1 (*60–62*) **(Fig. 6J and fig.S8L)**, a stark contrast compared to WT.

To determine whether loss of ATG5 Cys19 affects molecular pathways associated with aging, we performed proteomic analysis of aging hallmarks (**Fig. 6L)**. To do so we mapped proteins significantly changed in C19A livers onto a compendium of proteins with known roles in age-related pathologies (*75*). These results revealed a selective enrichment of aging-related pathways upon fasting in ATG5 C19A livers linked to canonical hallmarks of aging **(Fig. 6M)**. In particular, senescence and inflammation associated factors such as CD14 and CD9 were upregulated, indicating an enhanced inflammatory and SASP-like milieu. DNA repair and genomic instability markers including FEN1 and ERCC2, together with the chromatin associated stress responder HMGB2 and the pro-apoptotic factor BAK1, were elevated, consistent with increased DNA damage and stress signaling in mutant livers. Moreover, as observed by IHC **(Fig. 6J)**, autophagy adaptor SQSTM1/p62 accumulated, reflecting impaired proteostasis. Consistent with these molecular signatures, application of the transcriptomic aging clock developed by Tyshkovskiy et al. (*76*) to our proteomic data revealed that fasted ATG5 C19A mice liver exhibited a significantly increased predicted age compared with WT controls **(fig. S8P)**. Collectively, these coordinated changes support the notion that loss of ATG5 Cys19, and compromised autophagic induction, is sufficient to drive a pathogenic signature that manifests increased abundance in numerous molecular hallmarks of age-related tissue dysfunction **(Fig. 6M and table S5).**

## DISCUSSION

In this study, we establish OxiDR as a comprehensive, tissue-resolved atlas of the cysteine redox proteome regulated by DR. By quantitatively mapping oxidation states of thousands of cysteine residues in multiple mouse tissues, OxiDR extends beyond descriptive redox profiling and enables systematic identification of functionally relevant redox-sensitive nodes within protein networks. This resource thus offers a conceptual and experimental framework for dissecting how dietary cues are translated into redox-regulated signaling events that shape organismal physiology. The selectivity we observe, in which regulation of small set of functional cysteines against a largely stable bulk redox background, illustrates why a site-resolved atlas is required to distinguish targeted redox signaling from global shifts in redox tone. Bulk or compartment-level redox measurements would not have nominated ATG5 Cys19, because DR does not measurably change the aggregate oxidation state of most tissues. By quantifying reversible oxidation at the level of individual sites across tissues, ages, and diets, OxiDR provides the resolution needed to identify functionally consequential nodes, complementing gene- and protein-level atlases of aging and positioning cysteine redox state as an additional regulatory layer relevant to the hallmarks of aging framework. We anticipate that this resource will support hypothesis generation well beyond autophagy, particularly for the many tissue-specific regulated cysteines that remain functionally uncharacterized.

Leveraging this atlas, we identify ATG5 Cys19 as a diet-responsive redox switch that directly links DR and acute nutrient restriction to autophagy regulation. Mechanistically, DR selectively modulates the redox state of Cys19, thereby enhancing ATG5 interaction with ATG10 and promoting efficient formation of the ATG5-12 conjugated complex. A central mechanistic implication of our findings is that redox control of ATG5 Cys19 operates as a licensing step upstream of ATG12 conjugation, rather than by altering the catalytic chemistry of the conjugation machinery itself. The ATG12–ATG5 conjugate functions as an E3-like platform that specifies the site of LC3 lipidation (*52*), and its assembly depends on the E2-like enzyme ATG10, whose catalytic cysteine forms the thioester intermediate with ATG12 (*55, 56*). Because ATG10 Cys68 is distinct from this catalytic residue and ATG5 Cys19 lies in the N-terminal ubiquitin-like domain rather than the C-terminal domain engaged during canonical conjugation, we propose the Cys19–Cys68 disulfide as a redox-gated docking interaction that increases the efficiency of an otherwise constitutive enzymatic step.

Autophagy is widely recognized as a conserved determinant of longevity and metabolic fitness, and our analysis places ATG5 Cys19 at the intersection of redox regulation, autophagic competence, and metabolic resilience. Although ATG5 C19A mice appear grossly normal under basal conditions, they display pronounced vulnerability under metabolic stress, revealing a context-dependent requirement for this redox sensitive residue. These findings align with emerging models in which autophagy is dispensable for baseline homeostasis but becomes essential for adaptation to physiological stress. Genetic autophagy deficiency is compatible with embryonic development but not with the neonatal period, and autophagy is broadly required for the lifespan and healthspan benefits conferred by DR and other longevity interventions (*34, 44*). Our C19A mice reveal that this requirement can be localized to the redox regulation of a single residue: animals develop normally yet cannot tolerate acute nutrient withdrawal, phenocopying the fasting-intolerant, aging-accelerated state expected when adaptive autophagy fails. This is consistent with pharmacological and dietary autophagy inducers that act upstream of the core machinery, and suggests that the redox status of ATG5 Cys19 may represent a physiologically tunable set-point through which nutrient state is transduced into autophagic capacity.

In summary, we identify ATG5 Cys19 as a redox-regulated node that couples dietary restriction to autophagy dependent metabolic homeostasis and adaptive health resilience. By combining a systems level redox atlas with mechanistic and physiological analyses, our work reveals a previously unappreciated redox control point in the autophagy pathway and underscores the broader principle that precise cysteine redox regulation is a key determinant of organismal healthspan. Furthermore, several limitations should be acknowledged. The basis for the observed sex-specific liver phenotype warrants further investigation, potentially reflecting differences in hormonal regulation, redox buffering capacity, or metabolic wiring.

## MATERIALS AND METHODS

### Mouse lines, food and dietary restriction

Male C57BL/6 mice were obtained from the National Institute on Aging (NIA) aged rodent colony. All mice were individually housed in standard ventilated cages under a 12-hour light-dark cycle with controlled temperature and humidity. Mice were maintained on either an *ad libitum* fed (AL) or calorically restricted diet (dietary restriction, DR). AL fed mice had continuous access to standard chow (NIH-31), while CR mice were provided one pre-weighed pellet of NIH-31 fortified chow (3g) between 6:00 - 8:00 AM each day. The CR regimen was initiated at 14 weeks of age with a 10% reduction in caloric intake relative to AL controls. This restriction was increased to 25% at 15 weeks and then to 40% at 16 weeks, where it was maintained for the remainder of the study. Four cohorts of mice were included in the study: Young AL (4 months old at arrival to our facility), Young CR (4 months old at arrival to our facility), Old AL (18 months old at arrival to our facility), Old CR (18 months old at arrival to our facility). Upon arrival at the Harvard TH Chan School animal facility, all mice were individually housed and maintained under the same feeding conditions as described above for four weeks prior to tissue harvest (at 5 months or 19 months old, accordingly). All procedures involving animals were approved by the Harvard TH Chan School of Public Health Institutional Animal Care and Use Committee (IACUC) and were conducted in accordance with institutional and federal guidelines for the humane care and use of laboratory animals.

Fasting experiment was done using 6-month-old (24 weeks old) mice cohort, during the experiment all mice were individually housed. Only mice that are ≥25g for males and ≥20g for female will be subjected to fasting to ensure they have a good amount of fat to begin with at the start of the study. Humane endpoint of animal was determined as more than 20% of baseline body weight loss and/or marked lethargy (reluctance to move, unresponsive to gentle stimulation and/or Body condition score = 2). Mice were not re-used in any experiment until they recovered to their pre-fasting weight. All procedures involving animals were approved by the DFCI Institutional Animal Care and Use Committee (IACUC) and were conducted in accordance with institutional and federal guidelines for the humane care and use of laboratory animals.

### Maintenance of cell lines

HEK293T cells were cultured in DMEM (Corning, 10-017-CV) without pyruvate, supplemented with 10% FBS (GeminiBio, 100-106) and 1% P/S (Corning, 30-002-CI). AML12 (ATCC, CRL-2254) cells were cultured in DMEM/F12+GlutaMAX (Gibco, 10565-018), supplemented with 10% FBS (GeminiBio, 100-106), 1x Insulin-Transferrin-Selenium (Life Technologies, 41400045), 40ng/ml dexamethasone (Sigma-Aldrich, D4902) and 1% P/S (final: 100 I.U./ml Penicillin and 100 μg/ml Streptomycin; Corning, 30-002-CI). All cells were washed with PBS (Corning, 21-040-CV) detached using 0.25% trypsin (Gibco, 25200-056) and subcultured every other day.

### CPT synthesis

Succinimydyl iodoacetate (SIA) was purchased from Combi-Blocks and 6-aminohexylphosphonic acid hydrochloride salt (6-AHP) was purchased from SiKÉMIA. Synthesis was initiated by adding 6-AHP to SIA to final concentrations of 45 mM SIA and 175 mM 6-AHP, and allowed to react for 1 h at room temperature in the dark with gentle shaking. Reaction was quenched by addition of TFA to a final pH < 2, and purified on a Waters HPLC system (C18 column, solvent A: water with 0.035% TFA, solvent B: acetonitrile (ACN) with 0.035% TFA, 100%–40% solvent A over a 60-min gradient at a flow rate of 40 mL/min). The elution containing cysteine-reactive phosphate tag with the 6-carbon linker (6C-CPT) was frozen in a dry ice-acetone mixture for 30 min and lyophilized. Quality of synthesis was monitored by subjecting purified CPT to a 10-min run on Luna-HILIC column (Phenomenex) using an UltiMate-3000 TPLRS LC coupled with Q-Exactive™ HF-X mass spectrometer (Thermo Fisher Scientific).

### Tissue extraction

In order to preserve *in vivo* cysteine oxidation status, mice were euthanized by rapid cervical dislocation, and all tissues were extracted in less than thirty seconds following euthanasia. Wollenberger tongs were prechilled in liquid nitrogen and used to freeze-clamp each tissue immediately upon extraction(*77*).

### Protein CPT labeling from tissue extract

Following freeze-clamping, each tissue was homogenized in ice-cold 20% TCA using TissueLyser II (QIAGEN). The lysate from each biological replicate was split into two identical half-samples containing approximately 200 μg protein each then washed with 20% TCA, 10% TCA, and 5% TCA twice. One half-sample was resuspended in blocking buffer (100 mM HEPES pH 8.5, 2% SDS, 1 mM EDTA, 1 mM DTPA, 10 μM neocuproine, and 35 mM IAA) for 2 h at 37°C in the dark on a shaking incubator (1,200 rpm) to block all unmodified cysteine residues, while the other half sample was treated with labeling buffer (100 mM HEPES, 2% SDS, 1 mM EDTA, 1 mM DTPA, 10 μM neocuproine, and 35 mM CPT). After labeling, proteins in both half-samples were precipitated by methanol and chloroform, and resuspended in labeling buffer plus 5 mM TCEP to label reversibly modified cysteines.

### Protein digestion and TMT labeling for CPT-labeled cysteine proteome

Precipitated protein pellets were shortly airdried to evaporate methanol and resuspended in 100 μl 200 mM N-(2-Hydroxyethyl)piperazine-N′-(3-propanesulfonic acid) (EPPS) (pH 8.0), containing trypsin (Promega; final 1/100 enzyme/protein ratio) and LysC (Wako, Japan; final 1/100 enzyme/protein ratio). Protein lysates were digested overnight at 37 °C. Samples were centrifuged for 15 min at 15,000 x g, soluble peptide suspension was transferred into fresh Eppendorf tubes and peptide concentration was determined by BCA (Thermo Fisher Scientific). Equal amounts of peptides for each sample (150-200 μg) were transferred into fresh tubes and samples were completed to a volume of 100 μl with 200 mM EPPS (pH 8.0). Peptides were labeled with TMTpro 16-plex reagents in 30% ACN/EPPS solution for an hour at room temperature. A ratio-check was performed by mixing 2 μL of peptides from each channel, desalting using a stage-tip, and analysis by LC-MS. Upon completion of the ratio check analysis, the labeling reaction was quenched by addition of 5 μl hydroxylamine (5% stock) and incubation for 15 min at RT. The remainder of samples were mixed according to the total peptide loading ratios obtained from the ratio-check, and a second ratio-check was performed using 1% of the mixed sample to calculate a ratio to calibrate pipetting errors computationally for data analysis. CPT-labeled cysteine peptides were not included in the ratio-check analysis. Pooled samples at equal amount according to the ratio check were diluted with 12 ml of 1% FA in H_2_O and subjected to gravity flow driven C18 solid-phase extraction (200 mg Sep-Pak, Waters) and subsequently lyophilized.

### Cysteine peptide enrichment

TMT labeled pooled peptides were resuspended in phosphatase buffer (50 mM HEPES, 100 mM NaCl, 1 mM MnCl_2_ (pH 7.5) and lambda phosphatase (Santa Cruz Biotechnologies) was added according to the manufacturer’s instructions. Peptides were dephosphorylated during 2 hrs at 30°C shaking at 500 rpm. Subsequently, the sample was acidified with 10% TFA to a pH of < 3.0 (∼60 μl), subjected to gravity flow driven C18 solid-phase extraction (200 mg Sep-Pak, Waters) and vacuum dried. CPT labeled cysteine peptides were purified using the High-select Fe-NTA phosphopeptide enrichment kit (Thermo Scientific) according to the manufacturer’s instructions. Following elution of enriched CPT-labeled cysteine peptides, the peptide suspension is acidified with 10% TFA to a pH of < 3.0 (∼25 μl), subjected to gravity flow driven C18 solid-phase extraction (50 mg Sep-Pak, Waters) and vacuum dried.

### Cysteine peptide fractionation by HPLC

Dried peptides were resuspended in 300 μl of high-performance liquid chromatography (HPLC) buffer A containing 5 mM ammonium bicarbonate pH 8.0, 5% ACN and centrifuged through a PTFE 0.2 μM filter (Merck). Enriched cysteine-containing peptides were then fractionated using a high pH reversed-phase peptide fractionation by an Agilent 1100 quaternary pump with a degasser and a photodiode array with Zorbax 300Extened C18 column (Agilent, 2.1x 250mm 3.5-Micron). A 50-min linear gradient in 13 - 43% buffer B (5 mM ammonium bicarbonate, 90% ACN, pH 8.0) at a flow rate of 0.25 ml/min, and eluates were collected into a 96-deep-well plate. Fractions were consolidated into 12 tubes and vacuum dried followed by peptide desalting (stage tip) and LC-MS/MS analysis.

### Tissue sample preparation for global proteomics

Harvested tissues were lysed by 2% SDS, 100 mM EPPS buffer containing protease and phosphatase inhibitors (Thermo Fisher Scientific). Protein concentration was measured using BCA assay, followed by reduction with 10 mM DTT, alkylation with 40 mM IAA. After performing the methanol chloroform precipitation, samples were reconstituted with 50mM ammonium bicarbonate buffer and subjected to trypsin digestion overnight as described above. Samples were desalted and then loaded onto Evotip (Evosep EV2003) by following the manufacturer’s instructions. In brief, Evotips were washed with 20uL Buffer B (acetonitrile with 0.1% formic acid), and centrifuged for one minute at 800 x g. Additional 20uL of buffer A was added onto the Evotips and centrifuged one minute at 800 x g which were then soaked in 2-propanol for 20 seconds and centrifuged one minute at 800 x g with 20ul Buffer A loaded onto the Evotip. 20uL of samples were loaded and centrifuged for one minute at 800 x g followed by another 20uL of buffer A and centrifuged for one minute at 800 x g. Finally, 100uL of buffer A was load onto the Evotip then centrifuged for 10 seconds at 800 x g.

### Peptide desalting (stage tip) for LC-MS/MS

Peptides were desalted prior to LC-MS analysis using solid phase extraction (stage tip). Briefly, C18 Octadecyl HD solid phase extraction disk (CDS Analytics, USA) was used to prepare stage tips in house. The matrix was activated with 100% ACN, followed by washes with 70% ACN, 1% FA and H_2_O, 1 % FA. Dried peptides were dissolved in H_2_O, 1 % FA and passed through the C18 matrix. Peptides were washed twice with H_2_O, 1 % FA and subsequently eluted into MS vials in two steps with 40% ACN, 1% FA and 70% ACN, 1% FA. Peptides were vacuum dried and stored at -80°C until analysis.

### LC-MS/MS parameters

An Orbitrap Eclipse Tribrid Mass Spectrometer (Thermo Fisher Scientific) coupled with an Easy-nLC 1200 (Thermo Fisher Scientific) was used for proteomics measurements. Of each fraction ∼3 μg of peptides dissolved in 5% ACN, 5% FA were loaded onto an in-house 100-μm capillary column packed with 35 cm of Accucore 150 resin (2.6 μm,150 Å). Alternatively, peptides were loaded on a PepMap EASY Spray 75-μm C18 column (Thermo Fisher Scientific, 250 mm, 2 μm,100 Å). Peptides were separated and analyzed using a 180-min gradient consisting of 2% - 23% ACN, 0.125% FA at 500 nl/min flow rate. Spray voltage was set to 2500 with 300°C ion transfer tube temperature. A FAIMS Pro or FAIMS Pro Duo (Thermo Fisher Scientific) device was used for field asymmetric waveform ion mobility spectrometry (FAIMS) separation of precursors(*78*), and the device was operated with default settings and multiple compensation voltages (−40V/-60V/-80V). Under each voltage, positive mode data-dependent acquisition mode was used for a mass range of m/z 400-1400 applying two second cycle time method. Resolution for MS1 was set at 120,000. Singly-charged ions were not further sequenced, and multiply-charged ions were selected and subjected to fragmentation with standard automatic gain control (AGC) and 35% normalized collisional energy (NCE) for MS2, with a dynamic exclusion window of 30s. Quantification of TMT reporter ions was performed using the multinotch SPS-MS3 method(*79*) to measure 10 SPS precursors with HCD at 45% NCE for MS3 at 50000 resolution of orbitrap for scan range of 100-500 m/z.

Global proteomics samples were subjected to LC-MS analysis using Evosep ONE LC (Evosep Biosystems) coupled to a timsTOF HT mass spectrometer (Bruker). Peptides were separated using a PepSep C18 Column (15 cm × 150 μm, particle size 1.5 μm, Bruker) and a 15 SPD method on Evosep One. CaptiveSpray was operated at 1600 V with dry gas flow 3 L/min at 180 °C. The mass spectrometer was operated under parallel accumulation serial fragmentation (PASEF) mode for data-independent acquisition (DIA-PASEF) using the following parameters: polarity positive, scan *m*/*z* range 100–1700, mobility (1/K_0_) range 0.60–1.60V⋅s/cm^2^, ramp time 75ms, accumulation time 75ms, number of MS2 ramps: 20, number of MS2 windows: 60, mass range: 350-1250Da. Linear mobility-dependent collision energy was applied with 20eV at 0.60 V⋅s/cm^2^ and 59 eV at 1.6 V⋅s/cm^2^.

### Database searching

For CPT-length/labeling optimization experiments, the Comet algorithm was used to search all MS/MS spectra against a database containing sequences of mouse (*Mus musculus*) proteins downloaded from UniProt (https://www.uniprot.org/, 2022). Reversed sequences were appended as decoys for FDR filtering, and common contaminant proteins (e.g., human keratins, trypsin) were included. The following parameters were used for the database search: 25 ppm precursor mass tolerance; 1.0 Da product ion mass tolerance; fully tryptic digestion; up to three missed cleavages; variable modifications: oxidation of methionine (+15.9949), CPT modification on cysteine residues (+221.08169) for CPT labeled cysteine proteome and STY phosphorylation (+79.9663) for phosphor-proteomics. The target-decoy method was employed to control the false discovery rate (FDR)(*80–82*). To distinguish correct and incorrect peptide identifications, linear discriminant analysis (LDA) was used to control peptide-level FDR to less than 0.5%, and minimum XCorr was set to 1.0 during filtering. Peptides shorter than or equal to six amino acids length were discarded.

Data analysis of DIA-PASEF raw data generated by timTOF HT were performed using DIA-NN (version 2.2.0, https://github.com/vdemichev/DiaNN), searched against a database containing sequences of mouse (*Mus musculus*) proteins downloaded from UniProt (https://www.uniprot.org/, 2025). The parameters for generating *in silico* spectral library and search were as follows; protease: trypsin/p, missed cleavage: 2, peptide length range: 7-45, precursor charge range: 1-4, precursor m/z range: 300-1800, fragment ion m/z range: 200-1800, N-terminal M excision, cysteine carbamidomethyl and methionine oxidation. Default MS1 and MS2 accuracy, enable match between runs (MBR), Generic scoring, proteotypicity: protein names (from fasta), quantification strategy: QuantUMS, and library generation: IDs, RT & IM profiling.

### Site localization

A ModScore was calculated for each cysteine site in order to evaluate the confidence of site localization(*83*). This algorithm examines the presence of MS/MS fragment ions unique to each cysteine site on the same peptide to evaluate whether the best site match is correct when comparing to the next best match. Sites with Modscore ≥ 13 (p ≤ 0.05) were considered to be confidently localized. For quantification, the dataset was initially exported into two separate files, one containing sites from peptides with only one site quantified, and the other categorized as “composite”. Two files were manually merged into one for downstream analysis. Since cysteine is a rare amino acid, most sites are single sites and unambiguously localized on the parent peptides.

### Calculation of % oxidation on cysteines

All TMT channels were calibrated according to the second ratio-check described above to ensure the same amount of peptides were loaded to each channel prior to enrichment, and that pipetting errors were corrected. For each site, TMT reporter ion signal-to-noise ratio (S/N) from the oxidized cysteine channel was divided by S/N from the fully TMT-labeled cysteine channel of the same protein to obtain the % reversible oxidation value, effectively controlling for proteins abundance change. Values from five biological replicates were used to calculate the average and standard error of the mean (SE). All single and composite sites are listed in the supplementary tables. Single sites quantified from high confidence peptides (sum S/N > 160, isolation specificity > 0.75) were used for population data analyses. Unless otherwise stated, we considered sites with percent oxidation changes ± 15% and p value ≤ 0.05 as “changed significantly.” For redox network analyses we define cysteines with % oxidation ≥ 20% as extensively/highly modified.

### Data analysis

All analyses were performed in Perseus version 1.6.15.0, Prism 10.0 or R version 4.3.0 unless otherwise noted. Protein subcellular locations were downloaded from Human Proteome Atlas based on data reported previously(*84*) with experimental evidence, and matched to the Oximouse dataset. Nucleoplasm, nuclear speckles, nuclear bodies, nuclear membrane, nucleus, nucleoli (fibrillar center), and nucleoli were consolidated into the nucleus annotation. To minimize localization ambiguity, only proteins with single primary subcellular location reported were used. Secreted proteins did not present in the aforementioned database, and therefore singly-located proteins in “vesicles” category, and non-localized proteins were further queried from the secreted protein database downloaded from UniProt to generate a secreted protein list. Type I and II transmembrane proteins were downloaded from UniProt.

Heatmaps highlighting tissue-specific redox modifications were generated using a preprocessing with K-mean clustering (10 maximal iterations to both rows and columns) and using Euclidean distance where young and old tissues were separately clustered.

Protein networks were downloaded from BioPlex 3.0 where all data were generated with evidence from affinity pull-down and MS experiments(*29*). The % oxidation value of the highest oxidized site on each protein was used to represent the maximum extent to which this protein was ROS-modified and was mapped onto the BioPlex 3.0 network. Network visualization was performed in Cytoscape 3.10.0(*85*). Proteins with at least one site over 20% modified are regarded as highly-oxidized proteins, and interactions between two highly-oxidized proteins in the same tissue are considered a concerted oxidation event. To investigate whether the extent of coordinated redox modification in a protein network is more ubiquitous or tissue-selective, each network was scored by the percent of its constituent proteins that contain at least one highly oxidized cysteine in the same tissue. Non-identified proteins in communities were conservatively assigned as not highly oxidized to minimize false positive results. Communities with no highly modified proteins in any tissues were labeled as non-redox regulated. In addition, proteins that were reported as redox-sensitive previously(*30–32*) were extracted, mapped onto the highly-oxidized proteins, and overlap was analyzed in BioPlex 3.0 network to identify potential redox-regulated networks conserved across model organisms.

Analysis of proximal amino acids was performed by extracting motifs (lengths of 9 or 13 amino acids) centered around each quantified cysteine site in the dataset. Motifs were separated into two groups based on whether the motif contains the amino acid being tested (central cysteine excluded). Average % oxidation values of the central cysteines in these two groups were calculated, and the difference between the averages was used to determine whether the proximal amino acid being tested has a negative or positive influence on % cysteine oxidation. To visualize motifs, the pLogo algorithm was used(*86*). Motifs with total length of 13 amino acids were analyzed. For each tissue, the foreground contains motifs with highly oxidized central cysteines, the background contains all motifs quantified. Foreground sequences were not subtracted from the background.

Gene Ontology enrichment analysis was done with using DAVID gene ontology (https://davidbioinformatics.nih.gov/)(87) and g:Profiler (https://biit.cs.ut.ee/gprofiler/gost)(88) and shared evolutionary patterns between ATG5 Cys19 and ATG10 Cys68 was done by extracting the ortholog of each protein from species that are mapped in orthologous matrix (Oma) database (https://omabrowser.org/)(89) and multiple sequence alignment was performed for each protein to generate the cysteine conservation matrix result of each protein with 3 following categories. 10; indicates the cysteine site is conserved compared to human ATG5 Cys19 or ATG10 Cys68. 5; indicates protein itself is conserved compared to human but not the cysteine site, and 0; indicates neither cysteine site nor protein is conserved. Based on this matrix we computed row-wise proportional agreement to identify variables that are matched between ATG5 Cys19 and ATG10 Cys 68. Additional weight was assigned for matches categorized as 10 to highlight significant agreements. Fisher’s exact test was performed to assess the probability of proportional agreement occurring by chance, ensuring the robustness of results. All these analyses were done in group-wise manner for each taxonomic level.

AlphaFold3 based structural prediction of human ATG5 and ATG10 Cys68 fragment was performed by web based AlphaFold server (https://alphafoldserver.com/)(90) with two entries of full-length amino acid sequence of human ATG5 and ATG10 Cys68 containing fragment (FQTCLPMEEAFE).

Transcriptional age (tAge) prediction analysis was performed using Transcriptomic Age Calculator Online (TACO, https://app.gladyshevlab.org/TACO/), using our proteomics data, all the protein names were converted to ensemble gene name format for analysis. And data set was uploaded to server for further analysis using mouse clock of multi-tissue, mouse with elastic net algorithm, WT ad libitum group as control, split by tissue and sex as a covariate.

### Cysteine point mutant cell line generation

The HiFi Cas9 expression plasmid was generated by introducing a R691A mutation into the Cas9 coding sequence in pET-Cas9-NLS-6xHis (Addgene plasmid # 62933; Vakulskas et al., 2018). HiFi Cas9 protein was expressed in *Rosetta™(DE3)pLysS* Competent Cells (Novagen) and purified as described elsewhere(*91*). The sgRNA was generated using *GeneArt Precision gRNA Synthesis Kit* (Thermo Fisher Scientific).

To generate HEK293T cells carrying C19A mutation in ATG5, 0.6 μg sgRNA targeting sequence TGATATAGCGTGAAACAAGT was incubated with 3 μg HiFi Cas9 protein for 10 minutes at room temperature and electroporated into 2×10^5^ HEK293T cells along with a ssDNA oligo (catagtatggttctgcttccctttcagttatctcatcctgatatagcgtgaaaGCagttggaattcgtccaaaccacacatctcgaagcacatctttgtcatctg) using Neon transfection system *Kit* (Thermo Fisher Scientific). Mutants were identified by Illumina MiSeq sequencing. Briefly, the library was prepared using two-step PCR approach.

The primers for Amplicon PCR are 5’-ACACTCTTTCCCTACACGACGCTCTTCCGATCTAGTTTGGCTTTGGTTGAAGGA and 5’-GTGACTGGAGTTCAGACGTGTGCTCTTCCGATCTAGGTCCCCTTTGCACACTTA. The primers for Index PCR are: 5’-AATGATACGGCGACCACCGAGATCTACACTCTTTCCCTACACGACGCTCTTCCGATCT and 5’-CAAGCAGAAGACGGCATACGAGAT(n)_8_ GTGACTGGAGTTCAGACGTGTGCT. Aliquots of PCR product from each sample were pooled and purified using Qiagen PCR purification kit. The library was paired-end sequenced on an Illumina MiSeq according to the standard protocols (Illumina).

### Cysteine point mutant mice generation

ATG5 C19A point mutant mice were generated using CRISPR-CAS12a mediated genome editing. All RNA handling procedures were performed under RNAse free conditions. CAS12a protein was diluted to 1μg/μL in freshly prepared microinjection buffer (0.1x Tris-EDTA, TE), and crRNA and ssDNA donor templates were diluted to working concentrations of 10μM. For ribonucleoprotein (RNP) complex assembly, crRNA (final 1.2μM) was incubated with Cas12a protein (Final 0.122μM) at 37°C for 10 minutes. The ssDNA donor template containing the C19A substitution was then added (final 0.3μM) and incubated briefly at room temperature. The injection mixture was centrifuged through a 0.1μM filter to remove particulates and prevent needle clogging. Approximately 1μL of the CRISPR mixture was loaded into injection needles prepared using a micropipette puller and microforge. Microinjection was performed in fertilized one-cell embryos, which were subsequently transferred into pseudopregnant recipient females. Founder mice were genotyped by PCR amplification of the targeted locus followed by Sanger sequencing. The primers for Amplicon PCR are 5’-CCACCTTTTCTTTAAGTGAAGT and 5’-TTTAGAACTCAAACAGGGTTCT. Correctly edited founders were bred to establish germline transmission, and heterozygous offspring were intercrossed to generate homozygous ATG5 C19A point mutant mice.

### Indirect calorimetry measurement via Promethion Core system

The Promethion Core CGF system (Sable, Croydon) was used to perform IC measurement. The system is equipped with a baseline container to measure the oxygen consumption (VO2) and carbon dioxide production (VCO2) and energy expenditure. Food and water intake, body weight, movement were monitored simultaneously.

### Grip strength test

To measure grip strength, computerized grip strength meter (Bio-GS4, BIOSEB, Vitrolles, France) was used. Each mouse was held gently by the base of the tail and lowered toward the grid mesh of the apparatus until its paws firmly grasped the pull bar. The mouse was then pulled horizontally backward along the axis of the sensor with a steady, smooth and continuous motion until its grip was released. The digital force transducer recorded the peak resistance force in Newtons. Each mouse underwent four consecutive trials with an inter trial rest interval of approximately 15 seconds to prevent muscle fatigue. The average of the four independent measurements were calculated and used for statistical analysis.

### Fluorescence Microscopy

8×10^4^ AML12 cells were plated per well of 4-well chambered cover glass, borosilicate glass 1.5 (Lab tek-II) cultured with DMEM/F12+GlutaMAX (Gibco, 10565-018), supplemented with 10% FBS (GeminiBio, 100-106), 1x Insulin-Transferrin-Selenium (Life Technologies, 41400045), 40ng/ml dexamethasone (Sigma-Aldrich, D4902) and cotransfected with pCDNA3-eGFP-ATG5 (WT, C19S and C19A) and pCDNA3-RFP-LC3B plasmids using lipofectamine2000. Media was changed to fresh media after 6 hours of transfection. Starvation condition cells were incubated 2 hours with EBSS buffer prior to analysis.

For autophagic flux assay using halo-tag technique, 2.5×10^4^ HEK293T cells were plated onto the 4-well chambered cover glass, borosilicate glass 1.5 (Lab tek-II) cultured with HEK293T cells were cultured in DMEM supplemented with 10% FBS. pMRX-HaloTag7-GFP-LC3B was transfected using lipofectamine 3000. Media was changed to fresh media after 6 hours of transfection. Prior to analysis cells were incubated with final 100 nM of tetramethylrhodamine (TMR) ligand and 20 nM of bafilomycin A1 for 2 hours then changed into fresh media.

Images of eGFP/RFP expressing cells were acquired with an Ti2-E motorized inverted microscope with Focus Plan Apo 100x oil system (Nikon). The 488 nm and 561 nm were used to excite eGFP and RFP respectively. For GFP/Halo-tag expressing cells were acquired with Zeiss 980 with 63× 1.4 NA Plan-Apochromat oil objective and Airyscan detector (Zeiss MicroImaging). The 488 nm and 561 nm were used to excite eGFP and MTR labeled Halo respectively.

### Co-Immunoprecipitation

Cell pellets transfected with pCDNA3-eGFP-ATG5(WT, C19S and C19A), pCDNA3-flag-ATG10 (WT, LD1, LD2 and C68A) and pCDNA-ATG12-myc were lysed with lysis/washing buffer containing PBS, 0.5% (v/v) IGEPAL and EDTA-free cOMPLETE protease inhibitor (Roche) then incubated in ice for 30 min with additional benzonase (Millipore) treatment. To capture eGFP and Flag-tagged proteins, GFP-Trap (ChromoTek) and M2-frag (Sigma) beads were used respectively. Beads were prewashed with lysis buffer and incubated with samples for 1.5 hours at 4°C. Following incubation, the beads were washed three times with washing buffer and subsequently eluted by incubating them with 2x LDS sample buffer (Invitrogen) containing 5% B-mercaptoethanol for 10 minutes at 60°C.

### Western blot

For western blot analysis, cultured cells were washed with PBS and harvested cell pellet was lysed in 2% SDS in PBS buffer supplemented with EDTA-free cOMPLETE protease inhibitor (Roche) then sonicated to shear chromatin using tip sonicator with 20% amplitude for 3 seconds (Qsonica). The protein concentration was determined using a Pierce™ BCA assay kit (Thermo Fisher Scientific, USA). Lysate was adjusted to equal protein concentration across samples and diluted with 4x NuPAGE LDS sample buffer (Thermo Fisher Scientific, USA) and samples were heated for 10 min at 60°C. Co-immunoprecipitation samples were directly loaded onto the gel. Samples were separated on 4-12 % NuPAGE BisTris (Thermo Scientific, USA) gels using either MOPS or MES SDS running buffer (Thermo Fisher Scientific, USA). Proteins were transferred to PVDF membranes using the iBLOT2 transfer system (Thermo Fisher Scientific, USA) with iBLOT2 PVDF transfer stacks (Thermo Fisher Scientific, USA). Membranes were blocked with 3% BSA (Sigma, USA) in TBS + 0.1% Tween (Boston BioProducts, USA). Membranes were incubated with primary antibodies overnight at 4°C. Membranes were washed three times with TBS + 0.1% Tween-20 (Boston BioProducts, USA) followed by incubation for 1 hour at room temperature with secondary antibodies were incubated with membrane with TBS + 0.1% Tween-20 (Boston BioProducts, USA) containing 3% BSA (Sigma, USA) buffer then washed three times with TBS + 0.1% Tween-20 (Boston BioProducts, USA).

### Hematoxylin and Eosin (H&E) staining and Immunohistochemistry (IHC)

All Tissue samples were submitted to the DFCI core Pathology Department for standard processing and FFPE block sectioning. 5μm sections were made and conventional H&E staining was done. Automated immunohistochemistry was performed using Leica BOND-RX (Leica Biosystems). Formalin-fixed, paraffin embedded tissue sections were subjected to onboard deparaffinization followed by heat induced epitope retrieval (HIER) using either BOND Epitope Retrieval Solution 1 (ER1: citrate-based, pH 6.0) or BOND Epitope Retrieval Solution 2 (ER2: EDTA-based, pH 9.0), as specified for each antibody. Primary antibodies were applied as follows: cleaved caspase-3 (Asp175, Cell Signaling Technology, #9664, 1:2000 dilution) following ER1 retrieval, phosphor-histone H2A.X (Ser139, 20E3, Cell Signaling Technology, #9718, 1:100 dilution) following ER2 retrieval and SQSTM1/p62 (Abcam, #ab91526, 10ug/ml) following ER1 retrieval. Signal detection was carried out using the BOND polymer refine detection kit (Leica Biosystems) according to the manufacturer’s instructions. Slides were counterstained, dehydrated and coverslipped using xylene based mounting medium. Scans were acquired using a Grundium Ocus microscope (Grundium) at 5x-40x and viewed with Aperio ImageScope software (ver12.4.6.5003).

### Quantification and Statistical Analysis

Data was processed and significance defined using the pipeline described in the section above. Data analysis was performed in Excel, R, Prism, and Pymol, as described above. All data (unless otherwise noted) were presented as mean ± SEM. P values were calculated using two-tailed Student’s t test for comparison of variables. Sites with percent oxidation changes ± 15% and p value ≤ 0.05. p value<0.05 are marked as “*”, p value<0.01 are marked as “**” and p value<0.001 are marked as “***”. Unless otherwise noted, all stated replicates are biological replicates.

## ACKNOWLEDGEMENTS

This work was supported by the Claudia Adams Barr Program, the Lavine Family Fund, the Pew Charitable Trust, NIH DK123095, NIH AG071966, the Mark Foundation, the Smith Family Foundation, and the American Federation for Aging Research to E.T.C. E.T.C. is an HHMI investigator. S.S. was supported by Basic Science Research Program through the National Research Foundation of Korea (NRF) funded by the Ministry of Education (RS-2024-00408095). H.X. was supported by NIH R00AG07346103 and the Rita Allen Foundation. We thank DFCI Pathology core, HMS Transgenic Mouse core and The Initiative for Genome Editing and Neurodegeneration in Department of Cell Biology at HMS (Jiuchun Zhang) for their support.

## AUTHOR CONTRIBUTIONS

S.S., W.M. and E.T.C. carried out study conceptualization, design and direct research. S.S. designed and performed experiments, analyzed data, interpreted results, and prepared figures. N.B assisted with cysteine conservation analysis. H.X. performed mouse DR experiments. X.Z. and J.Z. assisted with gene editing. E.K., Y.W., J.J.P., Y.L., M.S., M.A.S., assisted with mouse experiments. D.W. assisted with structural modeling. S.M.W. and S.S. built the website. M.C.P-M. and W.M. provided DR mice. C.L and H.M.C. assisted with experimental discussions. The manuscript was written by E.T.C. and S.S. with the help of all authors.

## SUPPLEMENTARY MATERIAL

**Figure S1.**
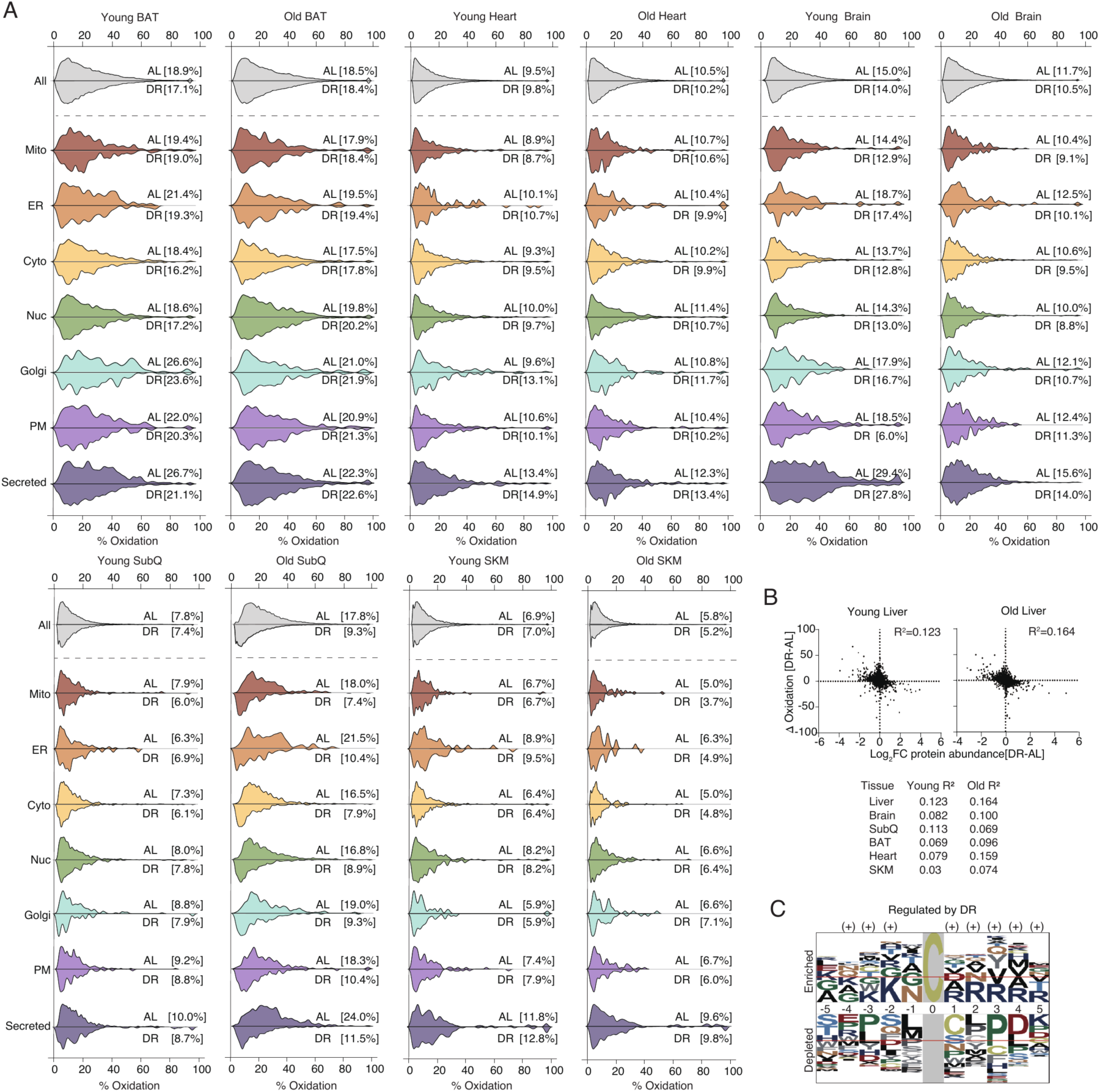
The *In vivo* cysteine oxidation landscape regulated by DR and characteristics of regulated cysteines. **A.** Distribution of cysteine percent oxidation across subcellular compartments in mouse tissues, corresponding to Figure 1E and 1F. Median percent oxidation values for each group are shown. **B.** Correlation analysis between protein abundance and the propensity for cysteine thiol oxidation in each tissue. n = 4 mice. **C.** Sequence motif analysis of significantly regulated cysteine sites.

**Figure S2.**
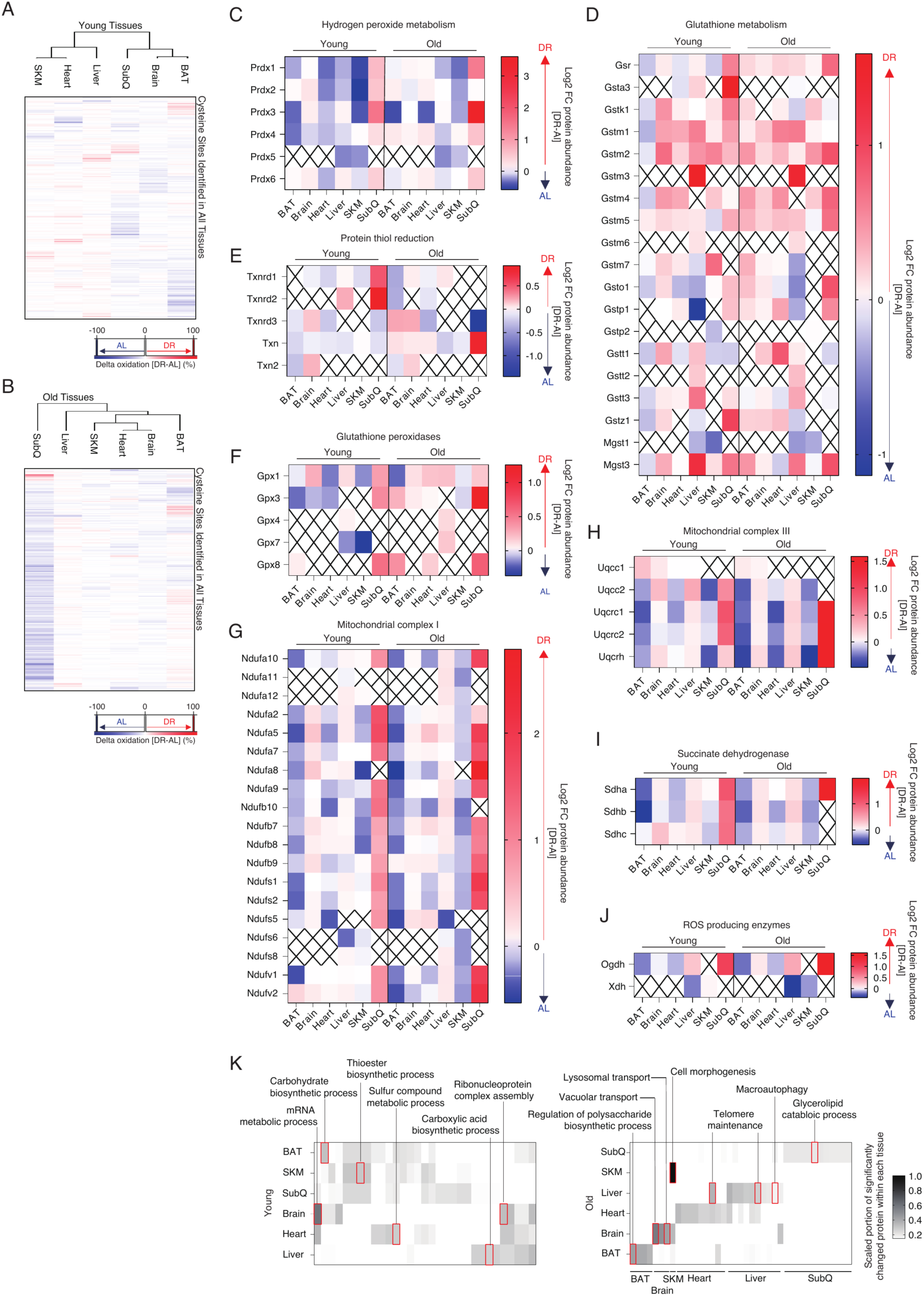
Tissue-specific profile of cysteine proteome and proteins controlling redox signaling networks. **A.** Heatmap cluster of cysteine sites showing concordant change in oxidation state upon DR across all tissues in young mice. Only the cysteine sites quantified in all tissues are presented. n = 4 mice per group. **B.** Heatmap cluster of cysteine sites showing concordant change in oxidation state upon DR across all tissues in old mice. Only the cysteine sites quantified in all tissues are presented. n = 4 mice per group. **C.** Tissue protein abundance measurements of proteins related to hydrogen peroxide metabolism comparing young and old AL and DR mice. n = 4 mice per group. **D.** Tissue protein abundance measurements of proteins related to glutathione metabolizing enzymes comparing young and old AL and DR mice. n = 4 mice per group. **E.** Tissue protein abundance measurements of proteins related to thiol reducing enzymes comparing young and old AL and DR mice. n = 4 mice per group. **F.** Tissue protein abundance measurements of proteins related to glutathione peroxidases comparing young and old AL and DR mice. n = 4 mice per group. **G.** Tissue protein abundance measurements of proteins related to mitochondrial complex I subunits comparing young and old AL and DR mice. n = 4 mice per group. **H.** Tissue protein abundance measurements of proteins related to mitochondrial complex III subunits comparing young and old AL and DR mice. n = 4 mice per group. **I.** Tissue protein abundance measurements of proteins related to succinate dehydrogenase subunits comparing young and old AL and DR mice. n = 4 mice per group. **J.** Tissue protein abundance measurements of proteins related to ROS producing enzymes comparing young and old AL and DR mice. n = 4 mice per group. **K.** Scaled portion of significantly altered cysteine oxidation sites between AL and DR in young and old mice tissues, associated with enriched GO term (biological processes).

**Figure S3.**
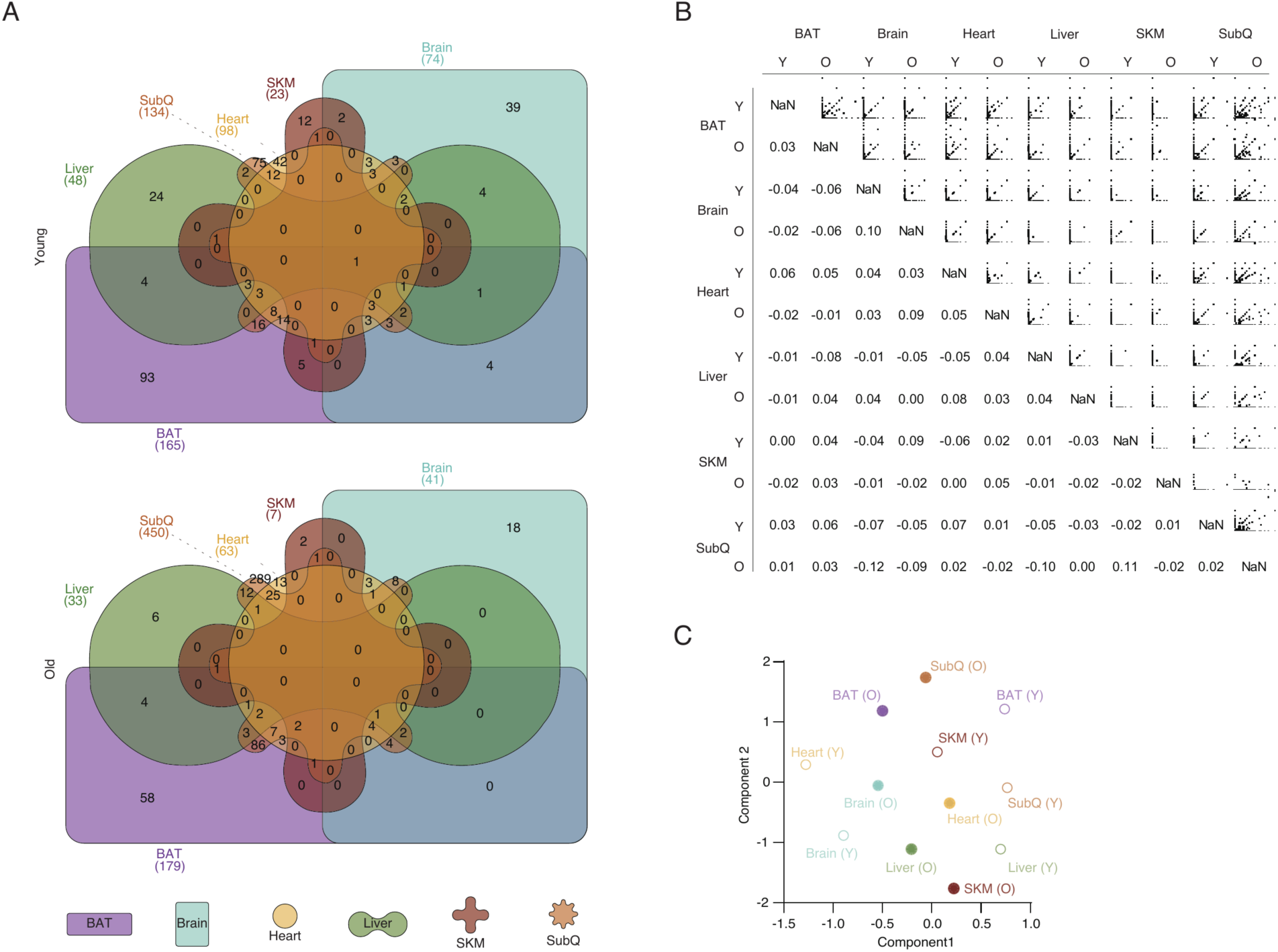
Systematic identification of redox regulated protein networks *in vivo.* **A.** Venn diagram of mapped redox regulated protein networks across tissues related to Figure 3D. **B.** Pearson’s correlation analysis of portion of mapped protein network by highly oxidized cysteine sites in each tissue. **C.** UMAP analysis of portion of mapped protein network by highly oxidized cysteine sites in each tissue.

**Figure S4.**
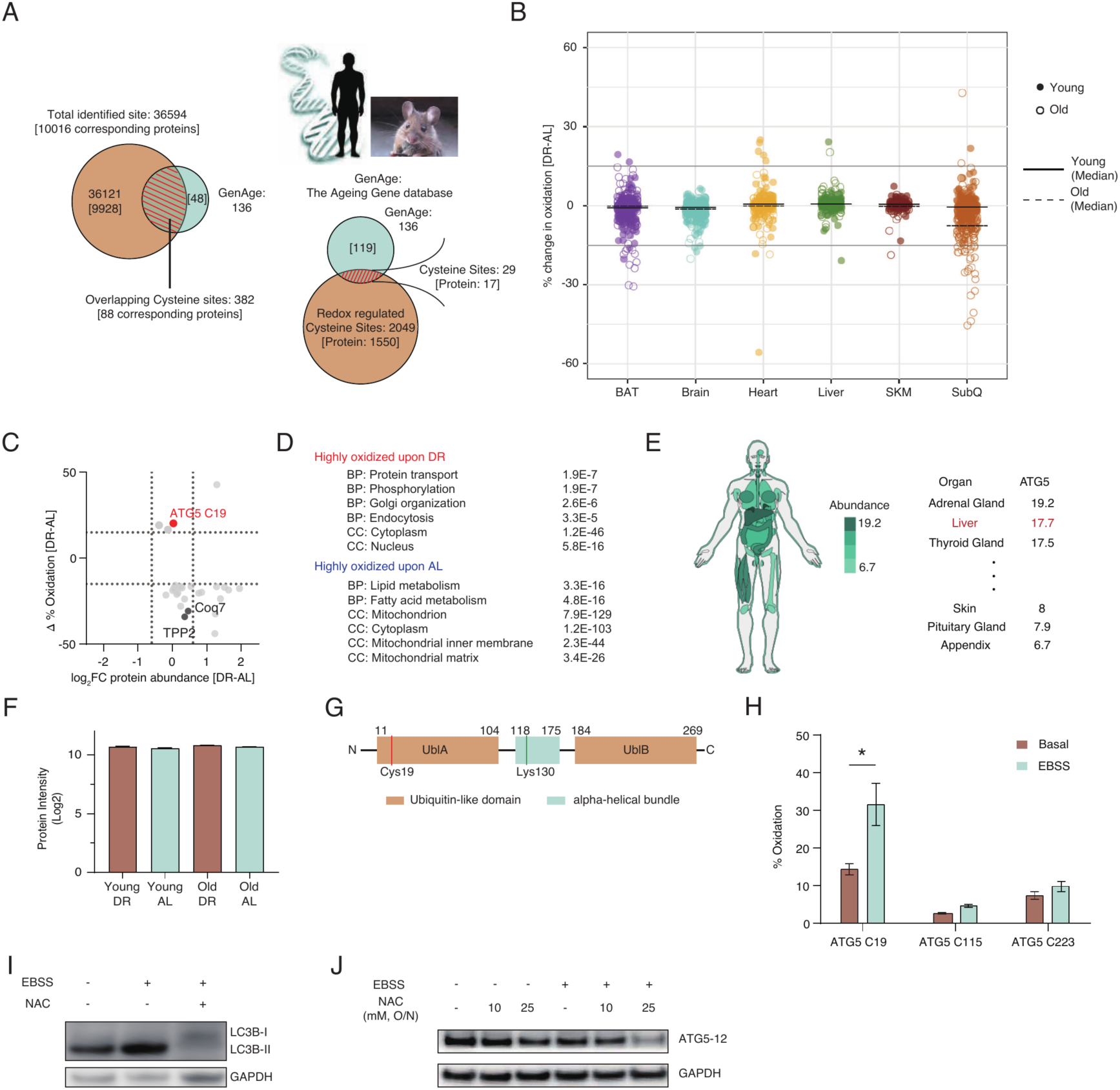
Selective oxidation of ATG5 Cys19 upon DR. **A.** Venn diagram comparing total proteins mapped by OxiDR (left) and proteins containing significantly redox-regulated cysteine sites upon DR (right) with the GenAge mouse database. **B.** Scatter plot showing percent changes in the oxidation of cysteine sites on corresponding proteins listed in the GenAge mouse database. Median percent change in oxidation values for each group are shown. **C.** Scatter plot of the significantly regulated cysteine sites upon DR shown in Figure 4A, plotted against their corresponding protein abundance changes upon DR. **D.** Gene ontology enrichment analysis of proteins containing highly regulated cysteine sites upon DR, related to Figure 4A. **E.** Abundance of ATG5 transcript in human tissues. **F.** ATG5 protein abundance in liver in young and old AL and DR mice. Data are presented as mean ± SD. n = 4 mice. **G.** Schematic of ATG5 functional domains. **H.** % reversible oxidation of individual ATG5 cysteine sites under basal and starvation (EBBS) conditions. n = 3 cell replicates per group. **I.** Immunoblot of LC3B following a 2hr treatment with 25 mM NAC, demonstrating reduced lipidated LC3B levels under starvation (EBSS) condition (EBSS treatment for 2 hrs). **J.** Immunoblot of ATG5-12 following overnight NAC treatment, showing gradual reduction of the ATG5-12 complex with increasing concentrations of NAC (Starvation/EBSS treatment for 2hr). p-value<0.05 is marked as “*”. Two-tailed Student’s t-test were used for pairwise comparisons. All bar plot data are presented as mean ± SD.

**Figure S5.**
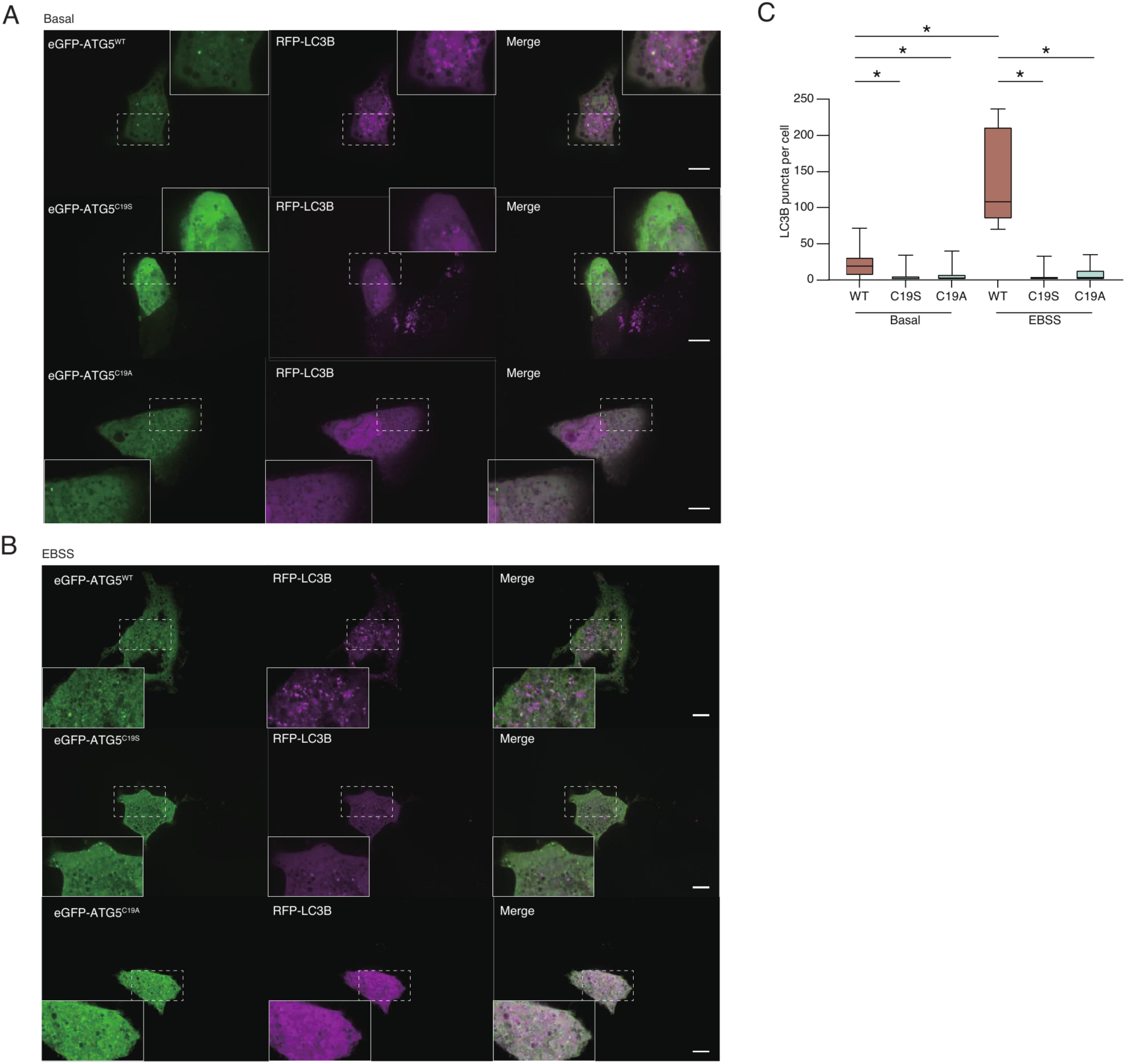
Effect of ATG5 Cys19 on LC3B puncta formation. **A.** Immunofluorescent images of GFP and LC3B in cells overexpressing GFP tagged WT ATG5, C19S ATG5, or C19A ATG5. scale bar = 10 µm **B.** Immunofluorescent images of GFP and LC3B in cells overexpressing GFP tagged WT ATG5, C19S ATG5, or C19A ATG5 after 2 hours EBSS. scale bar = 10 µm **C.** Quantitative analysis of LC3B puncta per cell, corresponding to the results shown in **(A)** and **(B)**. n = 7-39 cell replicates per group. p-value<0.05 is marked as “*”. Two-tailed Student’s t-test were used for pairwise comparisons. All bar plot data are presented as mean ± SD.

**Figure S6.**
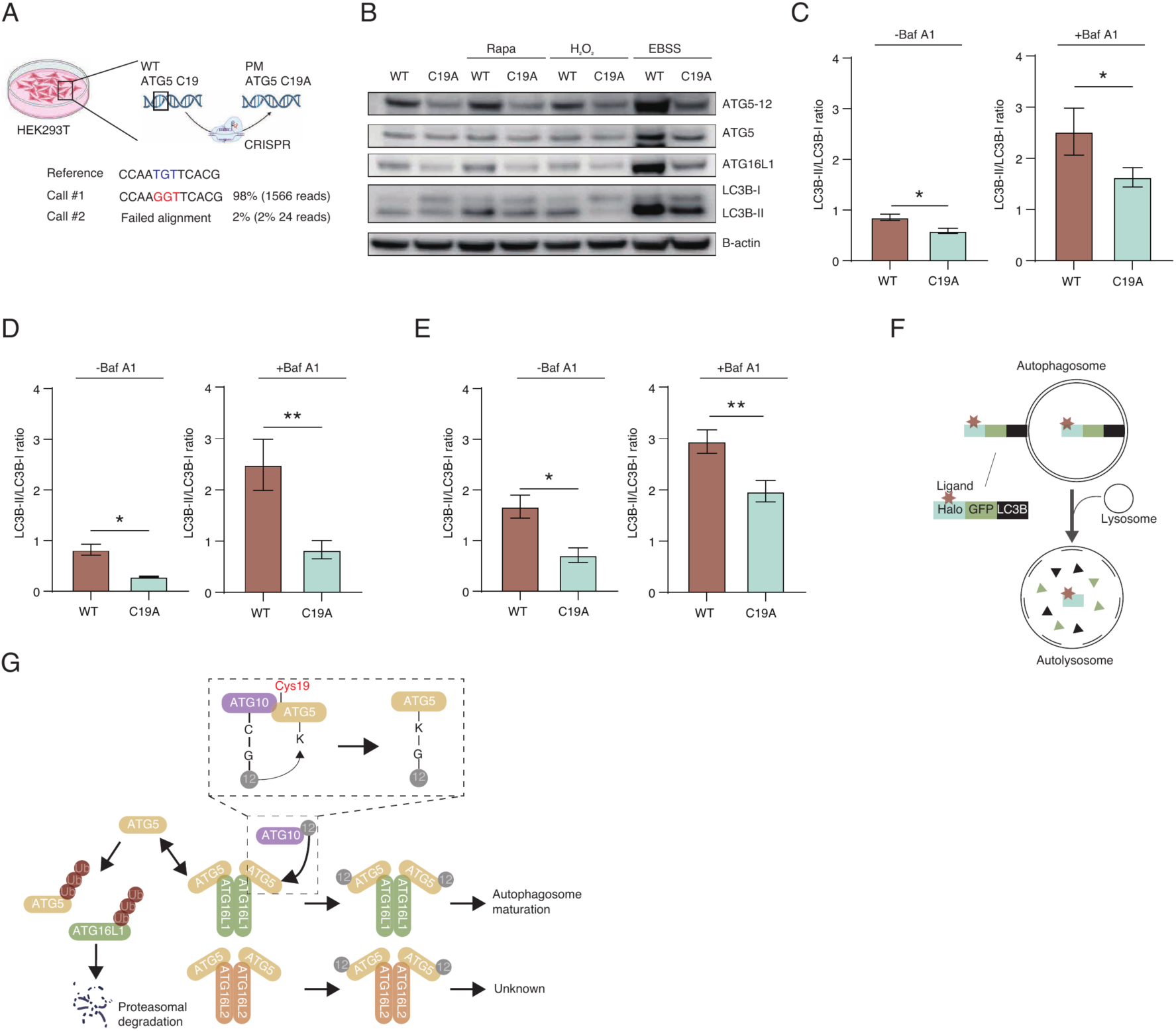
Role of ATG5 Cys19 on autophagy and ATG5-12-16L1 complex formation. **A.** Schematic of the endogenous ATG5 Cys19 point mutation using CRISPR and validation. **B.** Immunoblot of proteins relevant to autophagosome formation in WT and C19A ATG5 cells under various autophagy inducing conditions. **C.** Densitometric ratio of lipidated LC3B-II to non-lipidated LC3B-I measured by immunoblot with or without Bafilomycin A1 pretreatment (10 nM, 2hr) under rapamycin treated conditions. n = 2-3 cell replicates **D.** Densitometric ratio of lipidated LC3B-II to non-lipidated LC3B-I measured by immunoblot with or without Bafilomycin A1 pretreatment (10 nM, 2hr) under hydrogen peroxide treated conditions. n = 2-3 cell replicates **E.** Densitometric ratio of lipidated LC3B-II to non-lipidated LC3B-I measured by immunoblot with or without Bafilomycin A1 pretreatment (10 nM, 2hr) under EBSS treated conditions. n = 2-3 cell replicates **F.** Schematic of Halo-GFP-LC3B assay utilized to measure autophagic flux related to Figure 4G-I. **G.** Schematic of ATG5-12 covalent conjugation pathway. ATG12 conjugated ATG10 interacts with ATG5 to transfer ATG12 to ATG5 to form ATG5-12 complex which then stably form a complex with either ATG16L1 or ATG16L2, non-complex formed free ATG5 and ATG16L1 are known to be sensitive to degradation. p-value<0.05 is marked as “*” and p value<0.01 as “**”. Two-tailed Student’s t-test were used for pairwise comparisons. All bar plot data are presented as mean ± SD.

**Figure S7.**
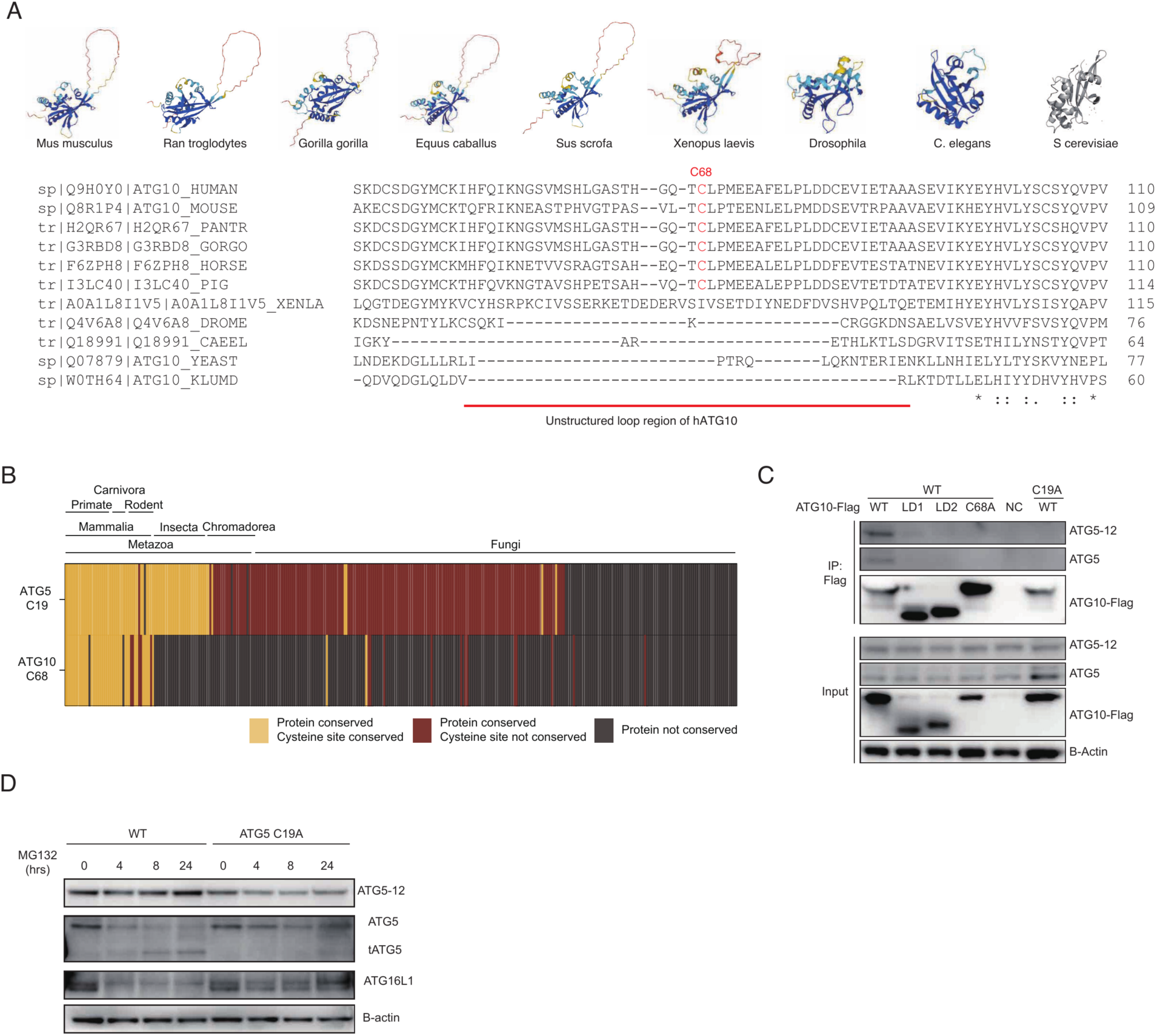
Evolutionary conservation and interaction between ATG5 Cys19 and ATG10 Cys68. **A.** Sequence alignment of ATG10 from various species, highlighting the evolutionary conservation of cysteine 68 (Cys68). **B.** Conservation analysis of ATG5 Cys19 and ATG10 Cys68 across diverse species in the OMA database, demonstrating a shared pattern conservation of co-conservation within *Mammalia*. **C.** Immunoprecipitation of ATG10-Flag constructs shown in Figure 5D in either WT or C19A ATG5 background cells. WT ATG5 exhibits higher affinity for ATG10. **D.** Immunoblot of WT and ATG5 C19A cells following treatment with the proteasome inhibitor MG132 (50 µM).

**Figure S8.**
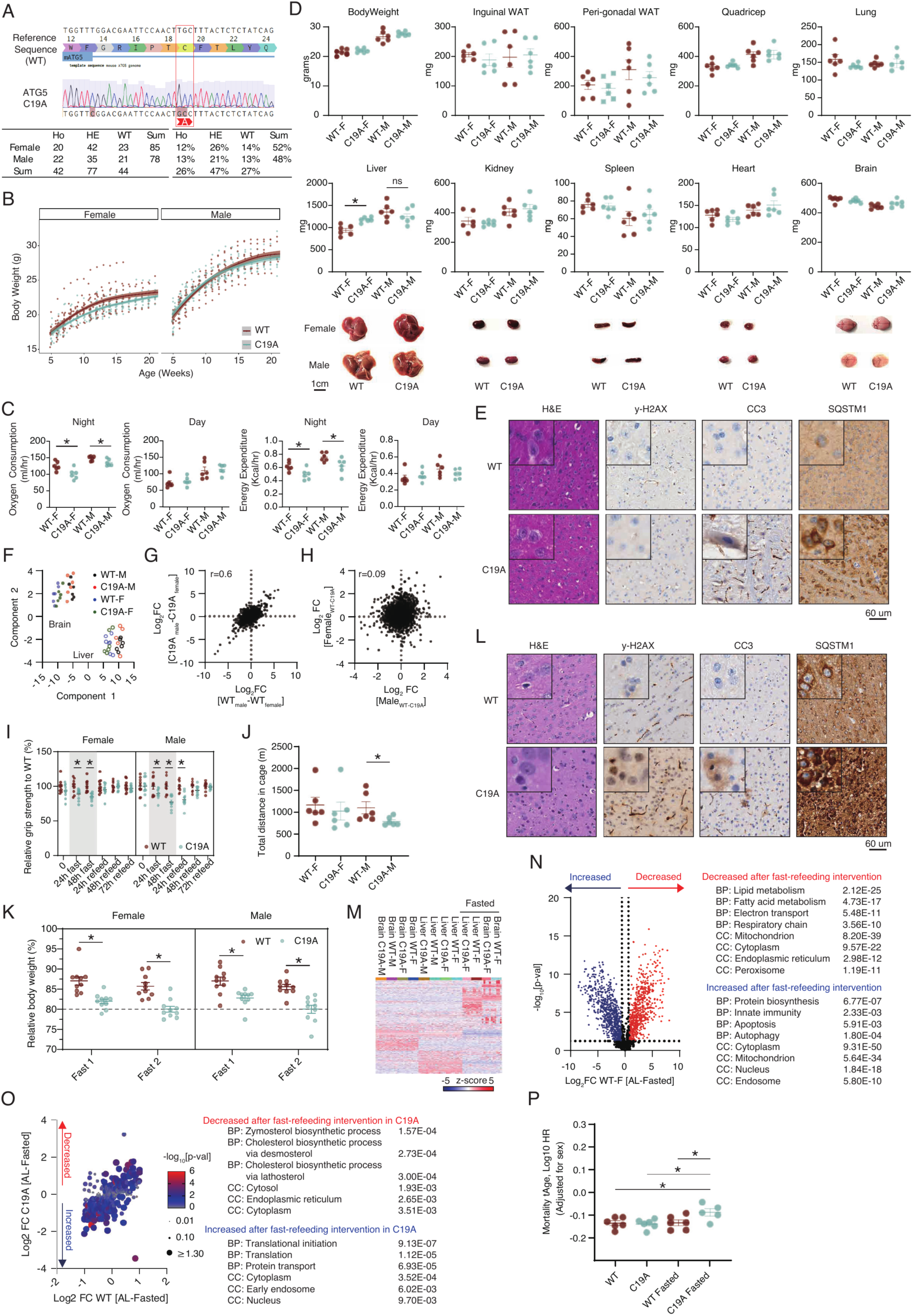
Systemic metabolic characterization of ATG5 C19A mice. **A.** Sequencing confirmation of the ATG5 C19A point mutation and Mendelian ratio of offspring. **B.** Body weight measurement of WT and ATG5 C19A mice during growth. n = 27-34 mice per group. **C.** Oxygen consumption (ml/hr) and energy expenditure (Kcal/hr) of WT and ATG5 C19A mice. n = 6 mice per group. **D.** Baseline measurements of bodyweight and tissue weights in female/male WT and ATG5 C19A mice. n = 6 mice per group. **E.** Hematoxylin and Eosin (H&E) staining and immunohistochemical (IHC) staining of y-H2AX, cleaved caspase-3 (CC3) and SQSTM1/p62 histological analysis of WT and ATG5 C19A mouse brain under AL feeding conditions. **F.** UMAP analysis of proteomic dataset derived from tissues of AL mice. n = 6 mice per group. **G.** Scatter plot of baseline protein abundance proteomics highlighting sex-dependent differences. n = 6 mice per group. **H.** Scatter plot of baseline protein abundance proteomics highlighting genotype-dependent differences across sexes. n = 6 mice per group. **I.** Grip strength measurement in female and male mice during the 48 hours fasting period and subsequent recovery. n = 10 mouse replicates. **J.** Total distance moved during 48 hours fasting and recovery. n = 6 mice per group. **K.** Body weight measurement after each 48 hour fasting cycle during the 2 week intermittent fasting intervention and normalized to the pre-fasting baseline of each cycle. n = 10 mice per group. **L.** Hematoxylin and Eosin (H&E) staining and immunohistochemical (IHC) staining of y-H2AX, cleaved caspase-3 (CC3) and SQSTM1/p62 histological analysis of WT and ATG5 C19A mouse brain following acute fasting and refeeding. **M.** Heatmap analysis of proteomics data at baseline and after the fast-refeeding intervention. n = 5-6 mice per group. **N.** Proteomic analysis and GO term enrichment analysis of differentially enriched proteins in liver upon fasting. n = 5-6 mice per group. **O.** Proteomic analysis of liver samples comparing AL and fasted conditions in both WT and ATG5 C19A mice that are significantly changed in ATG5 C19A liver only. n = 5-6 mice per group. **P.** Predicted mortality tAge differences between WT and ATG5 C19A liver under AL and fasted conditions. n = 5-6 mice per group. p-value<0.05 is marked as “*”. Two-tailed Student’s t-test were used for pairwise comparisons. All data are presented as mean ± SEM

**Table S1.** Summary of peptides, sites, and proteins identified/quantified from all mouse tissues.

**Table S2.** Cysteine site % oxidation values in tissues from young and old mice under AL and DR.

**Table S3.** The coordinated cysteine oxidation of BioPlex3.0 protein networks.

**Table S4.** Co-evolution analysis between ATG5 Cys19 and ATG10 Cys68 across different species.

**Table S5.** Proteomics analysis of WT and ATG5 Cys19A mice.

**Video S1.** WT (left) and ATG5 Cys19A (right) female mice after 48 hours of fasting, top view.

**Video S2.** WT (left) and ATG5 Cys19A (right) female mice after 48 hours of fasting, side view.

